# Infection-induced epilepsy is caused by increased expression of chondroitin sulfate proteoglycans in hippocampus and amygdala

**DOI:** 10.1101/2023.05.16.541066

**Authors:** Dipan C. Patel, Nathaniel Swift, Bhanu P. Tewari, Jack L. Browning, Courtney Prim, Lata Chaunsali, Ian Kimbrough, Michelle L. Olsen, Harald Sontheimer

## Abstract

Alterations in the extracellular matrix (ECM) are common in epilepsy, yet whether they are cause or consequence of disease is unknow. Using Theiler’s virus infection model of acquired epilepsy we find *de novo* expression of chondroitin sulfate proteoglycans (CSPGs), a major ECM component, in dentate gyrus (DG) and amygdala exclusively in mice with seizures. Preventing synthesis of CSPGs specifically in DG and amygdala by deletion of major CSPG aggrecan reduced seizure burden. Patch-clamp recordings from dentate granule cells (DGCs) revealed enhanced intrinsic and synaptic excitability in seizing mice that was normalized by aggrecan deletion. *In situ* experiments suggest that DGCs hyperexcitability results from negatively charged CSPGs increasing stationary cations (K^+^, Ca^2+^) on the membrane thereby depolarizing neurons, increasing their intrinsic and synaptic excitability. We show similar changes in CSPGs in pilocarpine-induced epilepsy suggesting enhanced CSPGs in the DG and amygdala may be a common ictogenic factor and novel therapeutic potential.

## Introduction

Epilepsy is a chronic neurological syndrome characterized by recurrent unprovoked seizures, associated with comorbidities affecting motor, sensory, cognitive, and psychological abilities. Seizures are a phenotypic correlate of neural network hyperexcitability resulting from aberrant firing of specific neurons in the brain. Genetic mutations in proteins critically involved in neurotransmission account for about 60% of epilepsy cases. The remaining 40% of patients have acquired epilepsy, resulting from brain insults that cause imbalance in excitatory and inhibitory neurotransmission that result in seizures. In the developed world brain injuries are the leading cause of acquired epilepsy, yet brain infection from various pathogens including viruses is a major contributor to epilepsy in the developing world. How a given insult leads to epilepsy, often referred to as “epileptogenesis”, remains poorly understood. Past studies have largely focused on molecular changes affecting the functions of neurons or glia cells^1–3^. However, given that brain injury and infection are associated with tissue remodeling, some recent studies have explored the role of extracellular matrix (ECM) in the etiology of epilepsy.

The ECM is a heterogeneous mixture of glycosylated proteins that fill the extracellular space (ECS) and constitutes about 20% of the brain volume^4^. ECM is broadly organized into three morphological features – basement membrane, perineuronal nets (PNNs), and neural interstitial matrix^5^. Basement membrane is a thin layer between blood endothelial cells and CNS parenchyma. It is mainly composed of collagen, laminin, fibronectin, integrins and dystroglycans. PNNs are dense lattice-like structures enwrapping neuronal soma and proximal dendrites comprising of proteoglycans, link proteins and tenascins. Interstitial matrix is an amorphous dense network of proteoglycans, hyaluronan, tenascins and link proteins. Proteoglycans are composed of a core protein with negatively charged glycosaminoglycan (GAG) side chains of repeating disaccharide units attached to it. The type of core protein and the carbohydrate composition of the GAG side chains determine the family of proteoglycan. Chondroitin sulfate proteoglycan (CSPG) is the most abundant proteoglycan found in the brain ECM.

In addition to being a structural scaffold for CNS parenchyma, ECM serves important functions in neurodevelopment, neural circuit formation, synaptic plasticity, learning and memory^6–8^. ECM also plays critical role during the recovery from brain injuries by supporting axonal growth and reorganization, myelination, synaptogenesis, and synaptic stabilization^9--11^. Furthermore, PNNs protect fast-spiking parvalbumin-containing (PV+) inhibitory interneurons against oxidative stress^12^ and support their high firing rate by decreasing membrane capacitance^13^. Owing to its highly negative charge, ECM is also proposed to function as an immobilized ion exchanger, and therefore, to regulate ionic and water homeostasis and the volume of ECS^14, 15^. Given these critical functions, disruption in the assembly and functions of ECM can contribute to neuronal network hyperexcitability and seizures. Indeed, varying changes in the components of ECM have been reported in patients with epilepsy and in animal models of epilepsy^16, 17^, however, it is not clear whether these changes are cause or consequences of seizures.

In the present study using a mouse model of Theiler’s murine encephalomyelitis virus (TMEV) infection-induced epilepsy, mice that develop epilepsy following infection showed a conspicuous increase in the expression of CSPGs in limbic areas of the brain, notably in the dentate gyrus (DG) and amygdala. The *de novo* expression of these CSPGs around excitatory neurons appears causal for the genesis of seizures as deletion of the acan gene, encoding for the most abundant CSPG aggrecan, in the DG and amygdala reduced seizure burden. Furthermore, patch-clamp recordings from dentate granule cells (DGCs) revealed an increase in intrinsic and synaptic excitability underlying TMEV infection-induced seizures. Cellular hyperexcitability was normalized in mice with aggrecan deletion in the DG and amygdala. Finally, we show that enhanced expression of highly negatively charged CSPGs around the DGCs causes the accumulation of K^+^, and potentially Ca^2+^, and increases their extracellular concentration, which in turn, depolarizes the resting membrane potential (RMP) of DGCs, increases synaptic excitability, and causes seizures. Increased CSPGs were also observed in other models of acquired epilepsy suggesting that CSPGs may be a novel more broadly applicable therapeutic target for acquired epilepsies.

## Results

### TMEV-infected mice with acute seizures show distinct changes in the expression of CSPGs in limbic areas

Previous studies have reported changes in the structure and composition of ECM associated with epilepsy in animal models and in the post-mortem brain tissues from patients with epilepsy^1, 18–20^. The most consistent finding has been a loss of PNNs around inhibitory PV+ interneurons in cortex and hippocampus^17, 21, 22^. Indeed, in a previous paper we showed that the release of matrix metalloproteinases (MMPs) from malignant gliomas degrade PNNs around PV+ interneurons thereby reducing their firing rate and causing tumor-associated epilepsy^13^. Viral brain infections are known to present with increased MMPs activity^23^, and hence, we wondered whether PNN disruption by MMPs may similarly explain viral infection-induced seizures. To answer this question, we infected C57BL/6J mice with TMEV, a well-established mouse model to study epileptogenesis following viral infection^24, 25^. We conducted immunohistochemistry (IHC) in brain slices obtained at 5 days post-infection (dpi) during peak acute seizure period (Fig. 1a) and visualized CSPGs using wisteria floribunda agglutinin (WFA), a lectin that selectively binds to N-acetylgalactosamine residues of CSPGs^26^. While a vast majority of TMEV-injected mice developed seizures, approximately 10-20% did not, allowing us to compare mice with seizures (TMEV S+) to mice without seizures (TMEV S−) to account for non-specific TMEV effects not pertinent to epileptogenesis.

**Fig. 1.**
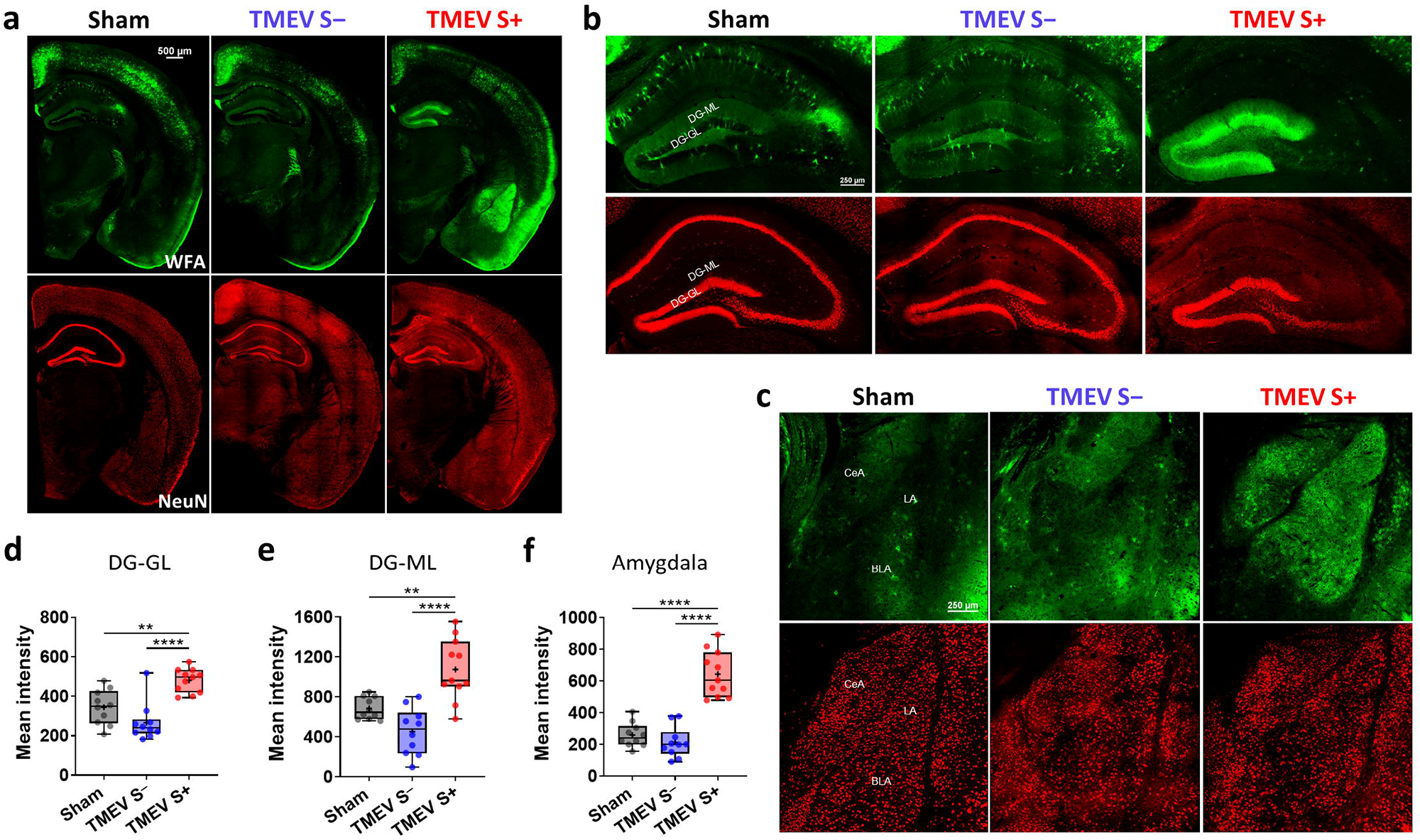
Increased deposition of CSPGs in dentate gyrus (DG) and amygdala of TMEV-infected mice with acute seizures. **a** Comparative images of the coronal brain hemislices obtained from mice treated with saline (Sham) or TMEV (TMEV S–: mice without acute seizures; TMEV S+ : mice with acute seizures) at 5 dpi and stained with the markers for CSPG (WFA, wisteria floribunda agglutinin, in green) and neuron (NeuN, in red). **b** Enlarged views of hippocampus show substantial increase in the level of CSPGs in the DG from the TMEV S+ group. A lack of CSPGs in the CA1 region is due to loss of CA1 pyramidal neurons common during acute TMEV-induced seizures. CSPG expression pattern and neuronal density in the TMEV S– group are comparable to the Sham group. GL, granular layer of DG; ML, molecular layer of DG. **c** Enlarged views of amygdala show substantial increase in the level of CSPGs in the TMEV S+ group. CSPG expression pattern in the TMEV S– group is comparable to the Sham group. CeA, central nucleus of amygdala; LA, lateral amygdala; BLA, basolateral amygdala. **d-f** Image analysis shows a significant increase in mean fluorescence intensity of WFA in the granular and molecular layers of DG (DG-GL (d) and DG-ML (e), respectively) and amygdala (f) from the TMEV S+ group compared to the Sham and the TMEV S– groups. Statistics: One-way ANOVA, Tukey’s multiple comparisons test; n=10-11 brain slices from 4-5 mice per group; **p<0.01, ****p<0.0001. Scale bar displayed in one image applies to all images in that panel.

Unexpectedly, rather than seeing a broad disruption of CSPGs throughout the hippocampus, we observed a striking increase in CSPGs expression in the granular and molecular layers of DG (Fig. 1b, d, e) and amygdala (Fig. 1c, f) only in the TMEV S+ group. We also observed a degradation of PNNs around inhibitory PV+ interneurons throughout the rest of hippocampus in TMEV S+ mice (Supplementary Fig. 1) except for area CA1 where TMEV-mediated loss of neurons (Fig. 1b) is well documented^25^. Representative examples show CA3 neurons with prominent degradation of CSPGs around the dentrites and to a lesser extent around somata (Supplementary Fig. 1a, b). Structual integrity analysis of PNN+ neurons shows disruption of the lattice-like PNN structure in mice with seizures compared to the control groups (Supplementary Fig. 1c-e, example from CA2).

MMPs are implicated in the proteolysis of ECM^27^ and increased activity of MMPs has been reported during epileptogenesis precipitated by tumors, traumatic brain injury (TBI), chemoconvusants, and other brain insults^27^. Gel zymography in whole hippocampal homogenate indeed showed increased enzymatic activity of MMP2 and MMP9 in the TMEV S+ group (Supplementary Fig. 2a, b, c). Furthermore, *in situ* zymography in hippocampal slices prepared from fresh frozen brains showed increased MMP2 and MMP9 activity, particularly pronounced in the CA1 and CA3 regions (Supplementary Fig. 2d, e). MMPs cleave core protein of aggrecan at specific sites thereby creating unique cleavage products of aggrecan. These expose new epitopes that can be identified using neoepitope-specific antibodies allowing one to directly measure the proteolytic activity of MMPs in vivo^28^. Western blots from whole hippocampal homogenates probed with aggrecan neoepitope antibody clone AF-28^29^ revealed aggrecan fragments (25-55 kDa) being markedly elevated only in the TMEV S+ mice compared to both control groups confirming MMP-mediated cleavage of aggrecan (Supplementary Fig. 2f-i).

Prior studies using mice either overexpressing or lacking MMP9 have implicated increased MMP9 activity in causing seiuzers following TBI and pentylenetetrazole-induced kindling^30, 31^. Hence, we injected MMP9^−/−^ mice with TMEV and monitored seizures by video and electrographic activity continuously from the DG-CA3 region of hippocampus up to 7 dpi. Typical electrographic activity corresponding to spontaneous behavioral seizure is shown in Supplementary Fig. 3a. All seizures identified are shown in the heatmap based on their severity score in Supplementary Fig. 3b. 88% (7/8) of MMP9^−/−^mice developed spontaneous seizures compared to 75% (6/8) of the wild-type (WT) mice, a difference that was not statistically significant (Supplementary Fig. 3c). Seizure frequency, severity, and duration were also comparable between MMP9^−/−^ and WT mice (Supplementary Fig. 3d-f). Comparisons based on seizure score found no difference between both the groups (Supplementary Fig. 3g, h) suggesting that increased activity of MMP9 alone is not sufficient to explain TMEV-induced seizures.

Since viral infection is likely inducing a number of MMPs, we tested minocycline, a broad-spectrum MMP inhibitor^32^ known to decrease TMEV-induced seizures^33^. Mice were treated with either minocycline (50 mg/kg) or vehicle twice daily during the first week of infection starting 2 hr before TMEV or sham injections (Fig. 2a). Seizures were observed a total of four times per day including during minocycline or vehicle treatment with an undisturbed period of at least 2 hr between two subsequent observations. Seizures were video-recorded to verify severity scores later. All seizures identified are shown in the heatmap based on their severity score (Fig. 2b). None of the sham-injected mice developed seizures (data not shown in the heatmap). The latency to first seizure and the percentage of mice that developed seizures (TMEV+vehicle – 12/12, 100%; TMEV+minocycline – 11/12, 92%) were not different between both the groups (Fig. 2c). However, minocycline treatment signficantly reduced average number of seizures during the entire acute seizure period (3-7 dpi) (Fig. 2d) and cumulative seizure burden, calculated as a cumulative sum of all seizure scores up to each dpi and used as a marker for seizure severity, between 5-7 dpi (Fig. 2e). Comparisons based on seizure score found significant reduction in severe seizures (stages 5-6) and mean seizure score (Fig. 2f, g). These results confirm an antiseizure effect of minocycline in the TMEV model. Mice were sacrificed at 7 dpi and IHC was conducted to measure the level of CPSG in the fixed brain slices. Mean fluorescence intensity of WFA staining was significantly increased in the granular and molecular layers of DG from TMEV-infected mice treated with vehicle compared to the sham-infected mice, and this increase was significantly reduced by minocycline treatment (Fig. 2h-j). However, minocycline did not completely normalize the CSPG level in DG as it was still significantly higher compared to the sham-infected mice. Similar results were obtained from amygdala as well (Fig. 2k, l). Minocycline treatment alone did not affect the expression of CSPGs in sham-infected mice. Minocycline treatment also reduced the formation of aggrecan fragments in the DG and amygdala resulting from augmented MMP activity in TMEV-infected mice (Supplementary Fig. 4). However, the decomposition of CSPGs in PNN found in areas CA3 and CA2 in mice with seizures was not noticeably different between minocycline and vehicle treatment groups. Taken together these results show a strong correlation between seizures, increased MMP activity, and the changes in CSPG levels during acute TMEV infection.

**Fig. 2.**
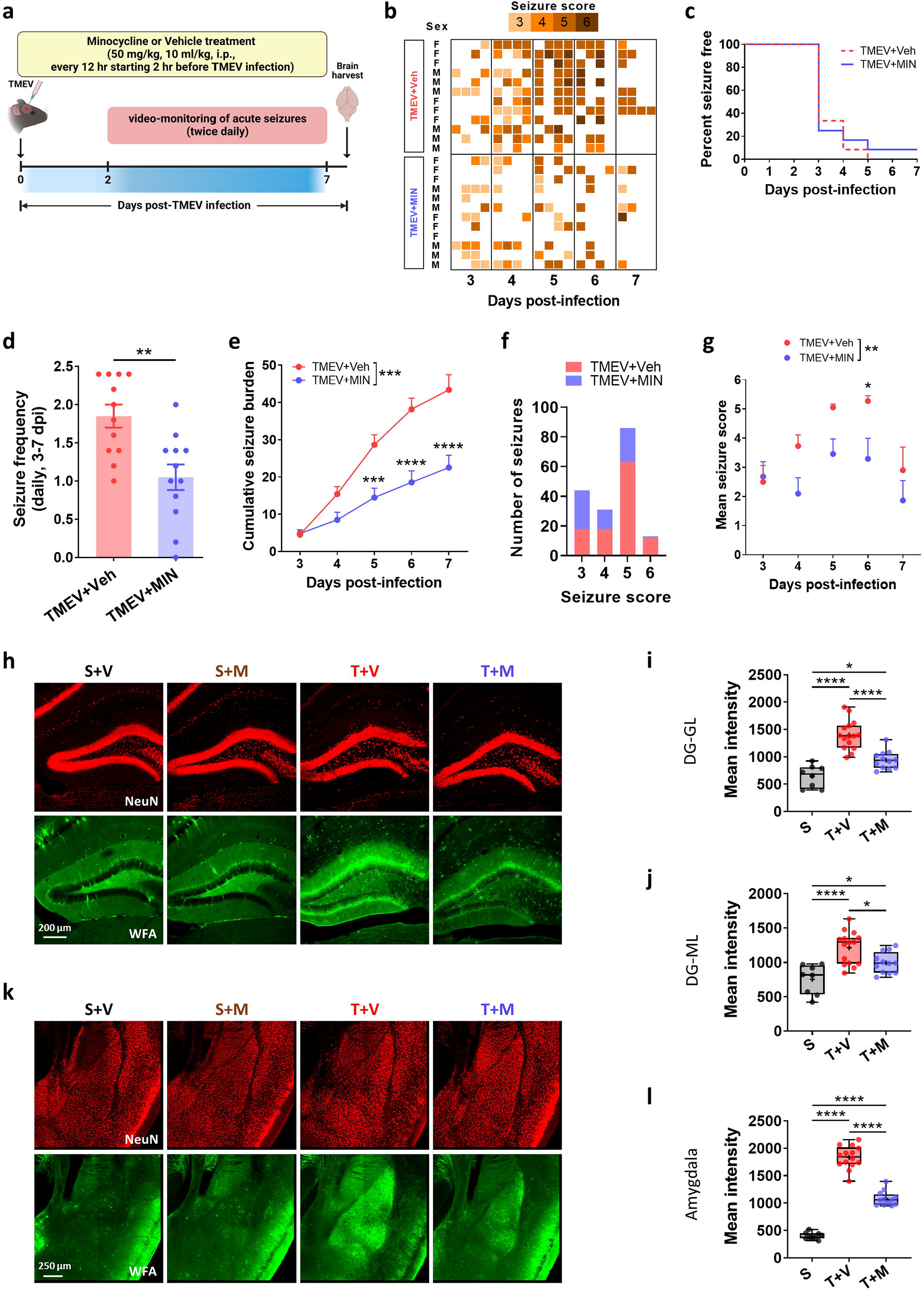
Minocycline treatment significantly reduces TMEV-induced acute seizures and inhibits the changes in expression of CSPGs associated with TMEV-induced acute seizures in DG and amygdala. **a** Experimental timeline. Mice were sacrificed at 7 dpi and the brains were either fixed for histology or were dissected into several regions and flash-frozen for biochemical analysis. **b** Heatmap shows handling-induced acute seizures observed based on their severity score for each mouse between 3-7 dpi. Mice were handled four times a day – during minocycline (MIN) or vehicle (Veh) injections twice daily and during seizure monitoring twice daily – with at least 2 hr of undisturbed period between each handling session. **c** Percentage of total infected mice in each group that remained seizure-free each day. None of the mice developed seizures before 3 dpi. **d** Average number of seizures per day between 3-7 dpi plotted for each mouse shows a significant reduction in seizure frequency by minocycline treatment (n=12, unpaired two-tailed t test, **p<0.01). **e** Average cumulative seizure burden, which is calculated as a mean of the summation of all seizure scores for each mouse up to each dpi, shows a significant reduction in seizure severity from 5 dpi by minocycline treatment (data shown as mean±SEM, n=12, two-way ANOVA, Šidák’s multiple comparisons test, ***p<0.01, ****p<0.0001). **f** Distribution of seizures based on seizure severity score. **g** Mean seizure score at each dpi shows significant reduction in TMEV-infected mice treated with minocycline (data shown as mean±SEM, n=12, two-way ANOVA, Šidák’s multiple comparisons test, *p<0.05). **h** Representative micrographs of the DG comparing the expression of CSPGs and neuronal density between mice infected intracortically with either TMEV (T) or Sham (S) and treated intraperitoneally with either minocycline (M) or vehicle (V). The brains slices were obtained at 7 dpi. **i-j** Image analysis shows that minocycline treatment significantly reduces an increased deposition of CSPGs (mean fluorescence intensity of WFA) in the granular and molecular layers of DG (DG-GL (b) and DG-ML (c), respectively) from TMEV-infected mice compared to the T+V group. The data from the S+V and S+M groups are combined as minocycline had no effect on the expression of CSPGs and neuronal density in mice infected with sham. **k** Representative micrographs of amygdala comparing the expression of CSPGs and neuronal density among four groups of mice. The brains slices were obtained at 7 dpi. **l** Similar to DG, minocycline treatment partially normalizes the expression of CSPGs in amygdala from TMEV-infected mice to the level in the sham group. Statistics (i-j, l): One-way ANOVA or Brown-Forsythe and Welch ANOVA, Tukey’s or Dunnett’s T3 multiple comparisons test; n=12-15 brain slices from 5-6 mice (T+V, T+M), n=8-9 brain slices from 2-3 mice (S+V, S+M); *p<0.05, ****p<0.0001. Scale bar displayed in one image applies to all images in that panel.

### Deletion of aggrecan in DGCs and amygdala significantly reduces TMEV-induced seizures

The next question was whether the dramatic increase in CSPG expression occurring in DG and amygdala during TMEV-induced acute seizures is a consequence or cause of seizures. If causal, suppression of CPSG synthesis prior to TMEV infection should eliminate seizures. To address this question, we deleted the *acan* gene that encodes for aggrecan specifically in the DG and/or amygdala neurons using *acan* conditional knockout mice (Acan^fl/fl^ mice) and cre recombinase. Aggrecan is a major CSPG constituent of both condensed PNN and the diffuse interstitial matrix in the brain. Genetic deletion of acan in neurons of Acan^fl/fl^ mice by cre-mediated recombination (either crossing Acan^fl/fl^ mice with cre-expressing mice or AAV9-mediated delivery of cre recombinase) has been shown to ablate aggrecan and prevent PNN formation^34^. For deletion of *acan* in the DG, we chose to cross Acan^fl/fl^ mice with POMC-cre mice. POMC (pro-opiomelanocortin-alpha) promoter drives expression of cre recombinase specifically in the granule cells of the DG, however, cre-mediated recombination starts to occur only at 2-3 weeks of age and remains spatially restricted until at least 24 weeks of age^35^. The expression of the target gene was found slightly decreased at 4 weeks, noticeably reduced at 12 weeks, and nearly absent at 16 weeks^35^. This lack of recombination in early developmental period is advantageous because CSPGs serve critical functions during development^8^, and therefore, deletion of aggrecan from birth is undesirable. Based on this earlier work and our pilot study to assess degradation of aggrecan in Acan^fl/fl^/POMC-cre^+^ mice, we used mice aged between 14-22 weeks for the TMEV-induced seizure study. For the deletion of aggrecan in neurons in amygdala, we injected an AAV9 vector construct driving cre expression under human synapsin promoter bilaterally in amygdala.

We generated three groups of mice – 1) Acan^+/+^ (*acan* WT), 2) DG-Acan^−/−^ (deletion of *acan* only in DGCs), and 3) DG-AMG-Acan^−/−^ (deletion of *acan* in DGCs and amygdala neurons) – and tested for their susceptibility to TMEV-induced acute seizures. The timeline for the study is shown in Fig. 3a. Our pilot work showed stable expression of cre recombinase and reporter gene and degradation of aggrecan in amygdala by 10 days of injecting AAV9 constructs. Therefore, we decided to infect these mice with TMEV 10-14 days post-AAV9 injections. Mice were video-monitored for seizures twice daily until 10 dpi. All seizures occurred between 3-8 dpi and are shown in the heatmap based on their severity score (Fig. 3b). Degradation of aggrecan had no adverse effect on normal mouse behavior, and the average weight of mice in each group was not different throughout the acute seizure period (Fig. 3c). Although all except one mouse in each group had at least one seizure by 8 dpi, 80% (8/10) of DG-AMG-Acan^−/−^ mice remained seizure-free until 4 dpi compared to 45% (5/11) of DG-Acan^−/−^ mice and 27% (3/11) of Acan^+/+^ mice showing increased latency to seizure development in DG-AMG-Acan^−/−^ mice (Fig. 3d). Average seizure frequency was significantly decreased in DG-AMG-Acan^−/−^ mice compared to Acan^+/+^ mice (Fig. 3e). Average seizure frequency was also reduced in DG-Acan^−/−^ mice compared to Acan^+/+^ mice, however, it was not statistically significant. Seizure severity was also decreased in DG-AMG-Acan^−/−^ mice compared to Acan^+/+^ mice as cumulative seizure burden showed significant reduction between 6-8 dpi (Fig. 3f). Cumulative seizure burden in DG-AMG-Acan^−/−^ mice was also significantly low compared to DG-Acan^−/−^ mice between 7-8 dpi. Comparisons based on seizure score found significant reduction in mean seizure score and severe seizures (stages 4-6) in DG-AMG-Acan^−/−^ mice (Fig. 3g, h).

**Fig. 3.**
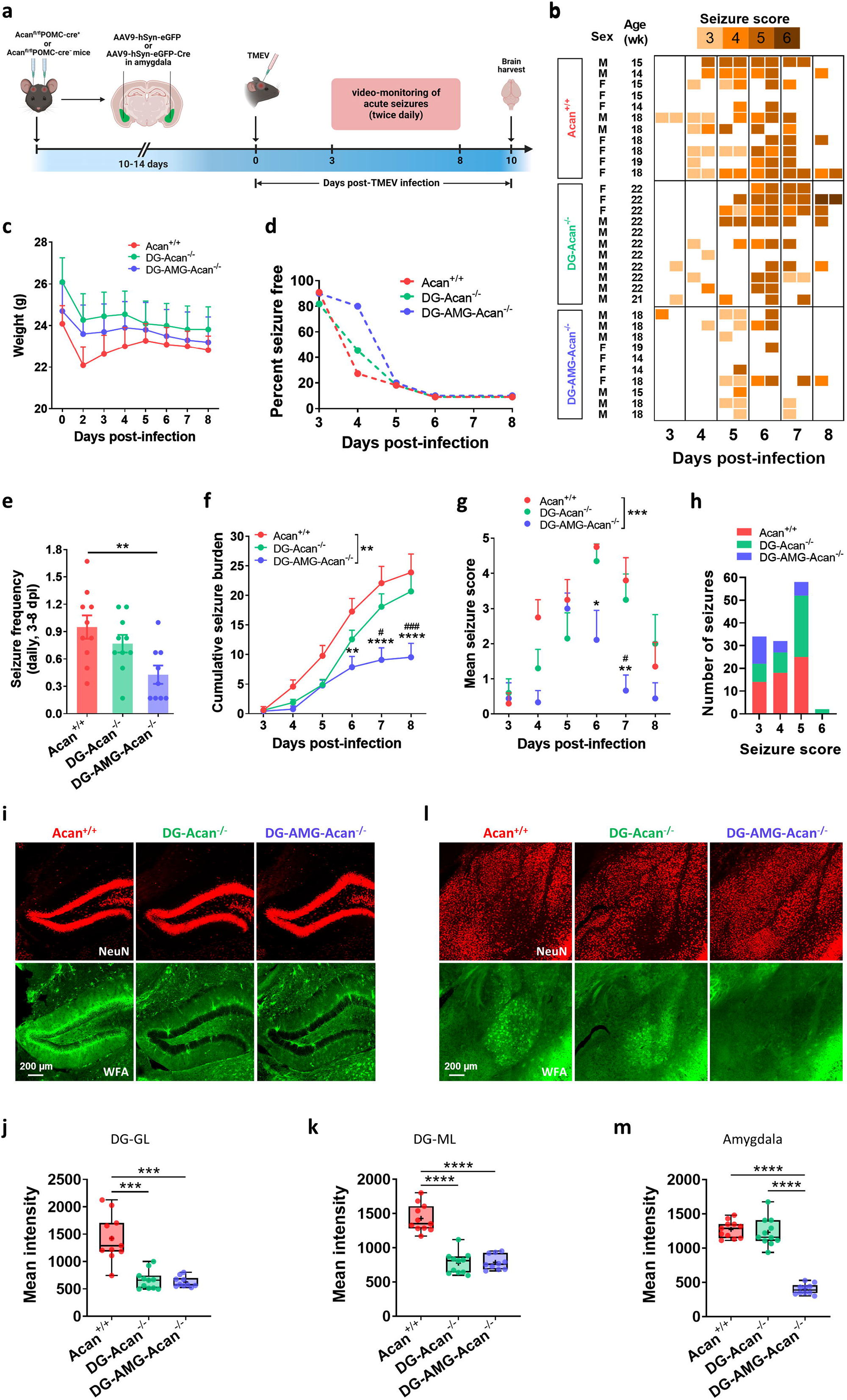
Deletion of aggrecan specifically in both DGCs and amygdala significantly reduces TMEV-induced acute seizures. **a** Experimental timeline. Acan^fl/fl^POMC-cre− mice were injected in amygdala bilaterally with AAV9-hSyn-eGFP to get Acan^+/+^ mice (control), whereas Acan^fl/fl^POMC-cre+ mice were injected in amygdala bilaterally with AAV9-hSyn-eGFP or AAV9-hSyn-eGFP-Cre to get DG-Acan^−/−^ and DG-AMG-Acan^−/−^ mice, respectively. All mice were infected with TMEV 10-14 days after injections of AAV9 constructs. **b** Heatmap shows handling-induced acute seizures observed based on their severity score for each mouse between 3-8 dpi. Seizures were induced twice daily by handling the mice with at least 2 hr of undisturbed period between each handling session. **c** Average weight of mice each day during acute TMEV infection period is not different for all three groups of mice. **d** Percentage of total infected mice in each group that remained seizure-free each day. None of the mice developed seizures before 3 dpi. **e** Average number of seizures per day between 3-8 dpi plotted for each mouse shows a significant reduction in seizure frequency in DG-AMG-Acan^−/−^ mice compared to Acan^+/+^ control mice (n=9-10, One-way ANOVA, Tukey’s multiple comparisons test, **p<0.01). **f** Average cumulative seizure burden, which is calculated as a mean of the summation of all seizure scores for each mouse up to each dpi, shows a significant reduction in seizure severity between 6-8 dpi in DG-AMG-Acan^−/−^ mice compared to the other groups (data shown as mean±SEM, n=9-10, two-way ANOVA, Bonferroni’s multiple comparisons test; comparisons between DG-AMG-Acan^−/−^ and Acan^+/+^ are denoted by *, whereas between DG-AMG-Acan^−/−^ and DG-Acan^−/−^ by ^#^; **p<0.01, ****p<0.0001, ^#^p<0.05, ^###^p<0.001). **g** Mean seizure score at each dpi shows significant reduction in DG-AMG-Acan^−/−^ mice (data shown as mean±SEM, n=9-10, two-way ANOVA, Šidák’s multiple comparisons test, *p<0.05, **p<0.01, ^#^p<0.05). **h** Distribution of seizures based on seizure severity score. **i** Representative micrographs show increased expression of CSPGs in DG at 10 dpi in Acan^+/+^ mice, but not in DG-Acan^−/−^ and DG-AMG-Acan^−/−^ mice, confirming cre recombinase-mediated deletion of acan gene in DGCs in DG-Acan^−/−^ and DG-AMG-Acan^−/−^ mice. **j-k** Image analysis shows significantly increased deposition of CSPGs (mean fluorescence intensity of WFA) in the granular and molecular layers of DG (DG-GL (b) and DG-ML (c), respectively) in Acan^+/+^ mice compared to DG-Acan^−/−^ and DG-AMG-Acan^−/−^ mice. **l** Representative micrographs of amygdala show increased expression of CSPGs at 10 dpi in Acan^+/+^ and DG-Acan^−/−^ mice, but not in DG-AMG-Acan^−/−^ mice, confirming cre recombinase-mediated deletion of acan gene in amygdala in DG-AMG-Acan^−/−^ mice. **m** Mean fluorescence intensity of WFA in amygdala is significantly higher in Acan^+/+^ and DG-Acan^−/−^ mice compared to DG-AMG-Acan^−/−^ mice. Statistics (j-k, m): One-way ANOVA or Brown-Forsythe and Welch ANOVA, Tukey’s or Dunnett’s T3 multiple comparisons test; n=10-11 brain slices (single slice per mouse); ***p<0.001, ****p<0.0001. Scale bar displayed in one image applies to all images in that panel.

To verify that *acan* deletion reduces expression of CSPGs in the DG and amygdala, mice were sacrificed at 10 dpi and IHC was conducted in the fixed brain slices. As in the TMEV-infected WT C57BL/6J mice with seizures (Fig. 1b), mean fluorescence intensity of WFA staining was significantly increased in the granular and molecular layers of DG from TMEV-infected Acan^+/+^ mice (Fig. 3i). Deletion of *acan* in DGCs indeed suppressed formation of aggrecan in TMEV-infected DG-Acan^−/−^ and DG-AMG-Acan^−/−^ mice (Fig. 3j, k), and delivering cre recombinase through AAV9 injections in amygdala significantly reduced formation of aggrecan in DG-AMG-Acan^−/−^ mice compared to sham AAV9-treated Acan^+/+^ and DG-Acan^−/−^ mice (Fig. 3l, m).

Taken together, inhibiting synthesis of aggrecan CSPG in both DGCs and amygdala neurons before TMEV infection significantly decreases frequency and severity of seizures. Although inhibiting synthesis of aggrecan only in DGCs achieves some reduction in seizures, it was not sufficient to make significant difference in overall seizure development compared to aggrecan WT mice. It is important to note that previous studies in the TMEV model of epilepsy have recorded severe electrographic activities from the DG-CA3 region of hippocampus and amygdala strongly implicating the interaction between these limbic regions in seizure generation^36, 37^. Therefore, it is concluded that increased deposition of CSPGs in the DG and amygdala is causally linked to increased seizure generation from TMEV infection.

### Upregulation of CSPGs in the DG and amygdala also occurs in pilocarpine-induced epilepsy

Given the involvement of hippocampus and amygdala in limbic seizure generation and the ictogenic role of enhanced deposition of CSPGs in the DG and amygdala in the TMEV model, we wondered if similar changes in CSPGs expression may occur in other models of temporal lobe epilepsy. Therefore, we measured CSPGs expression in the model of pilocarpine-induced status epilepticus (PISE). PISE is a widely used model of temporal lobe epilepsy triggered by the chemoconvulsant pilocarpine and present with the pathological changes and enhanced electrographic activities in several brain regions including hippocampus and amygdala^38^. Systemic administration of pilocarpine causes SE in mice and these mice develop spontaneous seizures rapidly within the first week of SE^39^. Supplementary Fig. 5a shows the WFA-stained brain slices comparing CSPGs expression between sham- and pilocarpine-treated mice at 5 days post-treatment. As in the TMEV model, CSPGs expression was found distinctly elevated in the DG granular and molecular layers as well as in amygdala from the pilocarpine-treated mice with SE compared to the control mice (Supplementary Fig. 5b-f). These results suggest that the increased deposition of CSPGs in the DG and amygdala in mice with limbic seizures is not restricted to a single model of temporal lobe epilepsy and it could be a pathological phenomenon contributing to seizures precipitated by various CNS insults.

### Increased excitability of DGCs during TMEV-induced acute seizures in WT C57BL/6J mice

We next asked how mechanistically increased deposition of CSPGs in the DG and amygdala may contribute to network hyperexcitability. We first focused on characterizing biophysical changes of individual DGCs by measuring their intrinsic electrophysiological parameters, firing rate, and synaptic currents during TMEV-induced acute seizure period. Patch-clamp recordings from DGCs in 300 µm thick acute horizontal brain slices were obtained between 4-10 dpi and the brain slices were fixed in 4% paraformaldehyde (PFA) solution after the recordings to measure the expression of CSPGs by IHC. Fig. 4a shows an example of WFA-stained brain slices confirming enhanced deposition of CSPGs in the DG from the TMEV group. RMP was significantly depolarized and input resistance was significantly increased in the TMEV-infected mice (Fig. 4b, c), whereas membrane capacitance was comparable between both the groups (Fig. 4d). Action potentials were recorded in current-clamp mode by injecting currents stepwise in 20 pA increment starting from -100 pA. Representative traces of action potentials corresponding to -100 pA and +100 pA current steps are shown in Fig. 4e. DGCs from mice with seizures generated a significantly higher number of action potentials induced by injecting currents in the range of 60-120 pA suggesting increased excitability of these cells (Fig. 4f). Voltage-clamp recordings were acquired as described in the methods to measure any changes in the properties of spontaneous as well as miniature excitatory and inhibitory postsynaptic currents (EPSC and IPSC) during seizures. Mean and cumulative distribution of amplitude and frequency of these recordings are compared in Fig. 4g-j. Mean amplitude and cumulative distribution of spontaneous EPSC (sEPSC) amplitude were significantly increased in the TMEV group (Fig. 4g). Although the mean frequency of sEPSC was unchanged, cumulative distribution of interevent interval of sEPSC was significantly decreased (Fig. 4g). In contrast, mean and cumulative distributions for amplitude and frequency of sIPSC were comparable between both the groups (Fig. 4h). For neural network independent miniature currents, mean amplitude and cumulative distributions of amplitude and frequency of both EPSC and IPSC were significantly increased in mice with seizures (Fig. 4i-j). In summary, DGCs from TMEV-infected mice with seizures exhibit depolarized RMP, increased firing rate, increased excitatory synaptic currents and increased miniature inhibitory synaptic current together tipping the excitatory-inhibitory balance into hyperexcitation contributing to seizure generation.

**Fig. 4.**
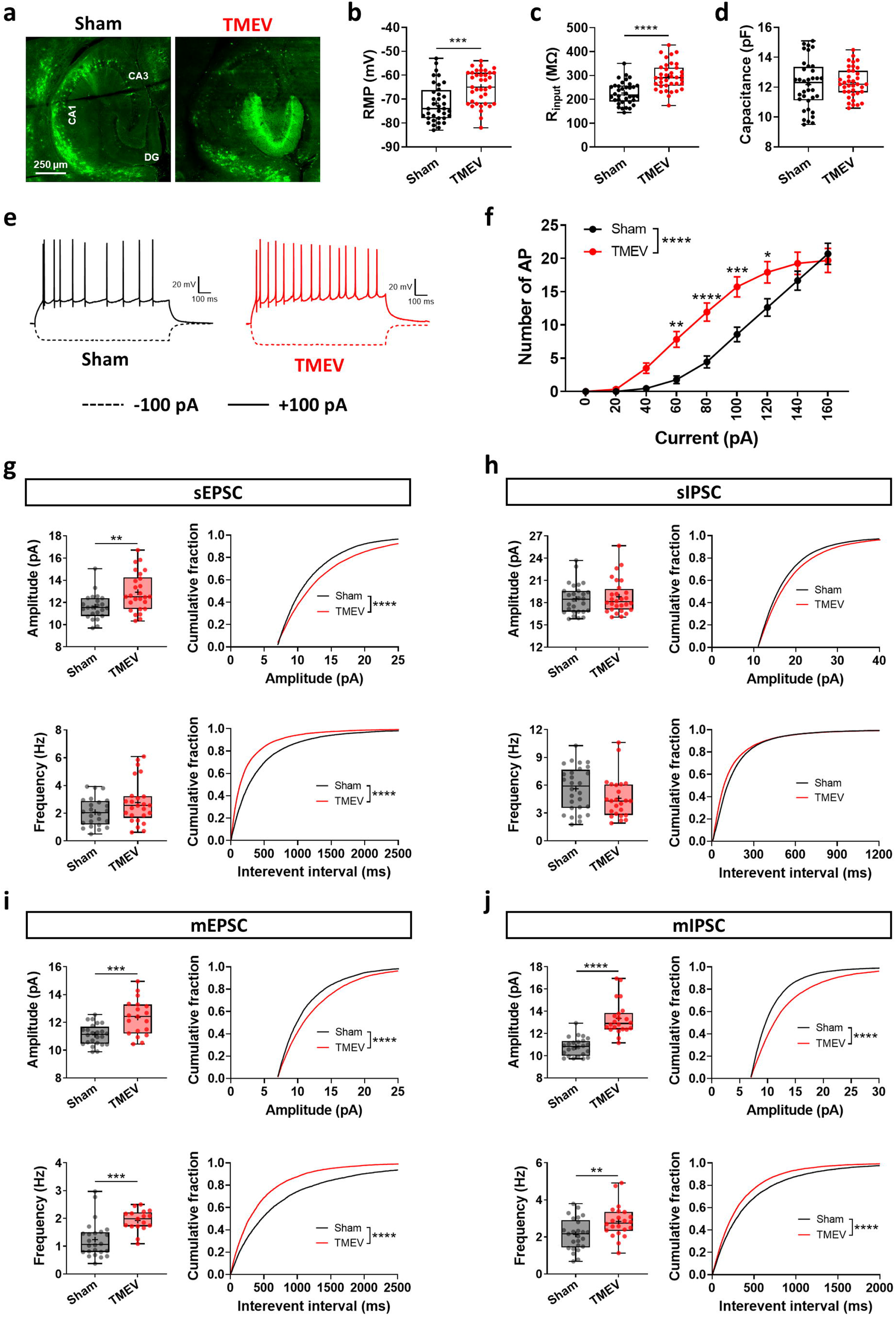
Increased intrinsic and synaptic excitability of DGCs during TMEV-induced acute seizures. **a** Representative micrographs of hippocampus from acute brain slices (300 µm thick) used for patch-clamp recordings show an increased level of CSPGs, as stained with WFA (green), in DG and degradation of CSPG in the CA1 region due to neuronal loss from TMEV-infected mice with acute seizures. The scale bar applies to both the images. **b** Mean resting membrane potential of DGCs is significantly depolarized in the TMEV group (n=36-37 cells from 4-6 mice, unpaired two-tailed t test, ***p<0.001). **c** Mean input resistance of DGCs is significantly increased in the TMEV group (n=36-37 cells from 4-6 mice, unpaired two-tailed t test, ****p<0.0001). **d** Mean membrane capacitance of DGCs shows no difference between the sham and TMEV groups. e Representative traces of action potentials recorded from DGCs of sham- and TMEV-infected mice. **f** Number of action potentials induced by injecting currents of varying strength in DGCs are significantly higher in TMEV-infected mice (data shown as mean±SEM, n=36-37 cells from 4-6 mice, two-way ANOVA, Šidák’s multiple comparisons test; *p<0.05, **p<0.01, ***p<0.001, ****p<0.0001). **g** A significant increase in mean amplitude (upper left), but not mean frequency (lower left), of sEPSC recorded from DGCs of TMEV-infected mice. Cumulative distributions show a shift toward higher amplitude (upper right) and lower interevent interval (lower right), indicating a higher frequency, of sEPSC from TMEV-infected mice. **h** No change in mean amplitude (upper left) and frequency (lower left) of sIPSC from TMEV-infected mice. Cumulative distributions of amplitude (upper right) and interevent interval (lower right) of sIPSC from TMEV-infected mice are comparable to the control mice. **i** A significant increase in mean amplitude (upper left) and mean frequency (lower left) of mEPSC from TMEV-infected mice. Cumulative distributions show a shift toward higher amplitude (upper right) and lower interevent interval (lower right), indicating a higher frequency, of mEPSC from TMEV-infected mice. **j** A significant increase in mean amplitude (upper left) and mean frequency (lower left) of mIPSC from TMEV-infected mice. Cumulative distributions show a shift toward higher amplitude (upper right) and lower interevent interval (lower right), indicating a higher frequency, of mIPSC from TMEV-infected mice. Statistics (g-j): bar graphs – n=23-28 cells from 4-6 mice (spontaneous), n=17-25 cells from 4-6 mice (miniature), unpaired two-tailed t test or Welch’s t test; cumulative fractions – Kolmogorov-Smirnov test; **p<0.01, ***p<0.001, ****p<0.0001.

### Genetic deletion of aggrecan rescues DGCs from TMEV-mediated hyperexcitability

To examine whether a decrease in TMEV-induced seizures observed in DG-AMG-Acan^−/−^ mice compared to Acan^+/+^ mice is also associated with reduced excitability of DGCs, we measured intrinsic electrophysiological parameters, firing rate, and synaptic currents by patch-clamping DGCs from these mice in acute horizontal brain slices between 4-10 dpi. The expression of CSPGs was verified in fixed brain slices following patch-clamp recordings. CSPG was indeed elevated in the DG from Acan^+/+^ mice, whereas a lack of CSPG enhancement was evident in the DG from DG-AMG-Acan^−/−^ mice (Fig. 5a). The input resistance and membrane capacitance were not altered but the RMP was significantly hyperpolarized in DG-AMG-Acan^−/−^ mice (Fig. 5b-d). Representative traces of action potentials corresponding to -100 pA and +100 pA current steps are shown in Fig. 5e. The input-output relationship of DGCs did not show any difference between both the groups (Fig. 5f). Interestingly, the RMP and the action potential firing rate of DGCs from Acan^+/+^ and DG-AMG-Acan^−/−^ mice were lower compared to WT C57BL/6J mice (Fig. 4b vs. Fig. 5d, Fig. 4f vs. Fig. 5f). The reasons for these physiological differences are not entirely clear but it could be explained by the difference in age of these mice during the patch-clamp studies (WT C57BL/6J mice – 8-10 weeks, Acan^+/+^ and DG-AMG-Acan^−/−^ mice – 22-24 weeks). To compare any other differences in action potential firing properties between TMEV-infected Acan^+/+^ and DG-AMG-Acan^−/−^ mice, action potential threshold current (minimum depolarizing current required to induce action potential consistently) was measured by injecting current gradually in small 2 pA steps. The representative traces of changes in voltage recorded corresponding to current injections steps are shown in Fig. 5g. The action potential firing threshold was found comparable between both the groups (Fig. 5h).

**Fig. 5.**
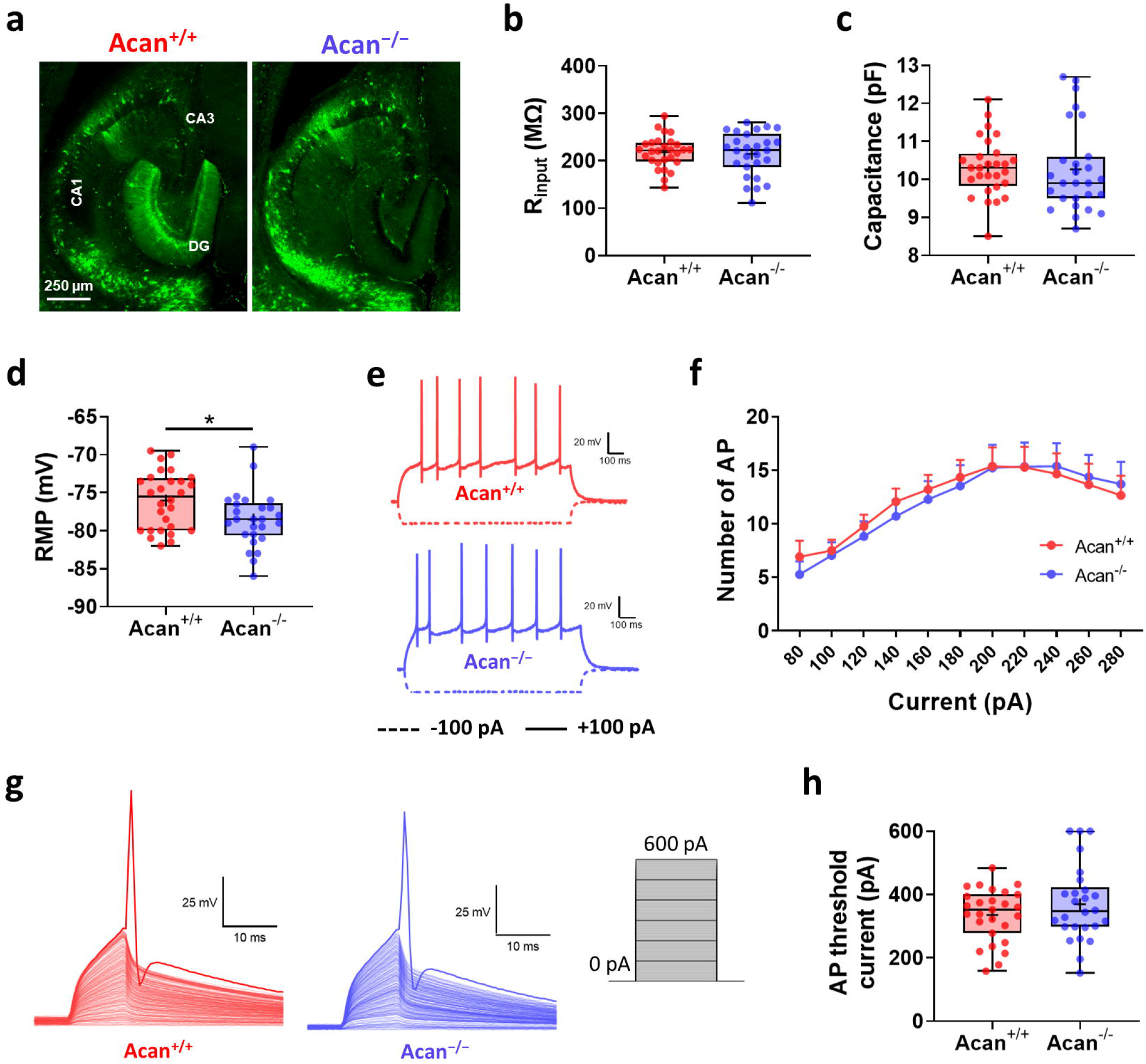
Resting membrane potential of DGCs from DG-AMG-Acan^−/−^ mice is hyperpolarized but no change in action potential firing rate and threshold current compared to Acan^+/+^ mice during acute TMEV infection. **a** Representative micrographs of hippocampus from acute brain slices (300 µm thick) used for patch-clamp recordings show a lack of increase in the level of CSPGs, as stained with WFA (green), in the DG from TMEV-infected DG-AMG-Acan^−/−^ mice with acute seizures. The scale bar applies to both the images. **b-c** Mean input resistance (b) and mean membrane capacitance (c) of DGCs show no difference between Acan^+/+^ and DG-AMG-Acan^−/−^ mice. **d** Mean resting membrane potential of the DGCs is significantly lower in DG-AMG-Acan^−/−^ mice (n=26-28 cells from 4-6 mice, unpaired two-tailed t test, *p<0.05). **e** Representative traces of action potentials recorded from DGCs from Acan^+/+^ and DG-AMG-Acan^−/−^ mice. **f** No difference in number of action potentials induced by injecting currents of varying strength in DGCs between Acan^+/+^ and DG-AMG-Acan^−/−^ mice. **g** Representative traces of voltage recordings in response to stepwise current injections from 0 to 600 pA (2 pA per step) to identify minimum amount of current needed to generate action potentials consistently (AP threshold current). **h** No difference in the AP threshold current between Acan^+/+^ and DG-AMG-Acan^−/−^ mice.

For spontaneous postsynaptic currents, there was no difference in mean amplitude and frequency of sEPSC, however, cumulative distributions showed a significant shift toward smaller amplitude and larger interevent interval in DG-AMG-Acan^−/−^ mice (Fig. 6a). In contrast, mean amplitude, but not cumulative distribution, of sIPSC was significantly increased in DG-AMG-Acan^−/−^ mice, whereas no difference was found in mean frequency and cumulative distribution of sIPSC between the groups (Fig. 6b). For miniature postsynaptic currents, amplitude of mEPSC and mIPSC were similar between both genotypes of mice (Fig. 6c, d). Whereas mean and cumulative distributions of mEPSC and mIPSC frequencies were significantly reduced in DG-AMG-Acan^−/−^ mice (Fig. 6c, d). Taken together, DGCs from TMEV-infected DG-AMG-Acan^−/−^ mice had hyperpolarized RMP and decreased excitatory synaptic currents which readily explains the decrease in seizures experienced by these mice compared to Acan^+/+^ mice. Although there was no change in the action potential firing rate and threshold current of DGCs from TMEV-infected DG-AMG-Acan^−/−^ mice, the overall results show that these cells receive less depolarizing input from the neuronal network, which in turn, reduces seizure generation.

**Fig. 6.**
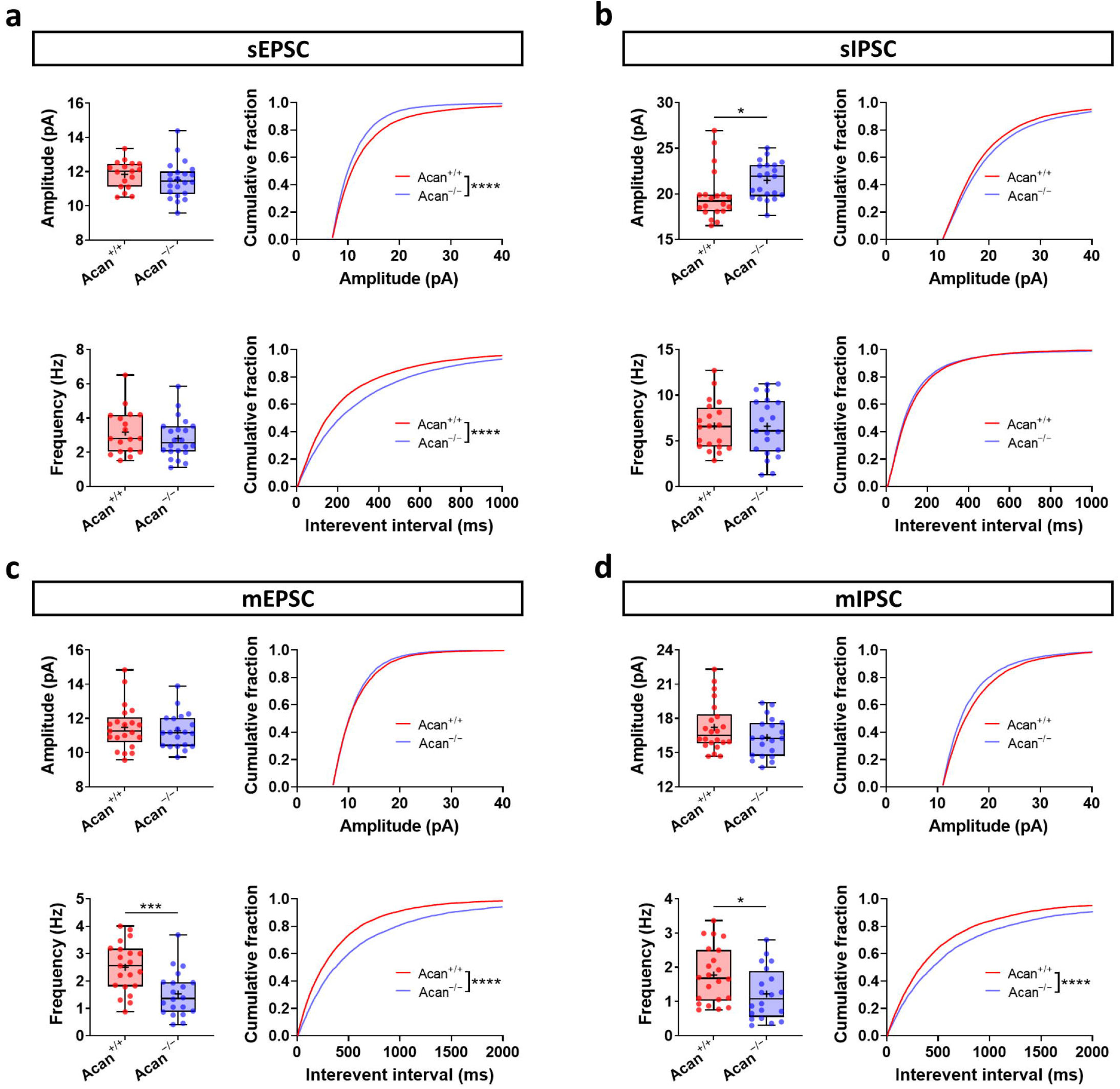
Decreased synaptic excitability of DGCs from DG-AMG-Acan^−/−^ mice compared to Acan^+/+^ mice during acute TMEV infection period. **a** No change in mean amplitude (upper left) and frequency (lower left) of sEPSC between Acan^+/+^ and DG-AMG-Acan^−/−^ mice. Cumulative distributions show a shift toward lower amplitude (upper right) and higher interevent interval (lower right), indicating a lower frequency, of sEPSC from DG-AMG-Acan^−/−^ mice. **b** A significant increase in mean amplitude (upper left), but not mean frequency (lower left), of sIPSC recorded from DGCs of TMEV-infected DG-AMG-Acan^−/−^ mice. No difference in cumulative distributions of amplitude (upper right) and interevent interval (lower right) of sIPSC between both groups of mice. **c** A significant decrease in mean frequency (lower left), but not mean amplitude (upper left), of mEPSC from TMEV-infected DG-AMG-Acan^−/−^ mice. Cumulative distributions similarly show a shift toward higher interevent interval (lower right), indicating a lower frequency, without any changes in amplitude (upper right) of mEPSC from DG-AMG-Acan^−/−^ mice. **d** A significant decrease in mean frequency (lower left), but not mean amplitude (upper left), of mIPSC from TMEV-infected DG-AMG-Acan^−/−^ mice. Cumulative distributions similarly show a shift toward higher interevent interval (lower right), indicating a lower frequency, without any changes in amplitude (upper right) of mIPSC from DG-AMG-Acan^−/−^ mice. Statistics (a-d): bar graphs – n=17-22 cells from 4-6 mice (spontaneous), n=20-22 cells from 4-6 mice (miniature), unpaired two-tailed t test; cumulative fractions – Kolmogorov-Smirnov test; *p<0.05, ***p<0.001, ****p<0.0001.

### No change in presynaptic density of excitatory and inhibitory terminals in the DG from mice with TMEV-induced seizures

Interestingly, the frequencies of sEPSC, mEPSC, and mIPSC recorded from the DGCs were increased in TMEV-infected WT C57BL6/J and Acan^+/+^ mice compared to their corresponding control mice. The frequency of synaptic currents is dependent upon the total number of presynaptic neuronal terminals that synapse onto the postsynaptic neuron and the quantal release probability at the individual synapses. Since CSPGs could affect synaptic organization^11^, increased frequency of presynaptic events in the DGCs from WT C57BL6/J and Acan^+/+^ mice might have resulted from an increased density of presynaptic boutons during acute TMEV infection. To test this hypothesis, we immunostained brain slices for vGLUT1 and vGAT, which are the markers of excitatory and inhibitory presynaptic terminals, respectively, at 5 dpi^40^. The expression of CSPGs and vGLUT1 are shown in the representative volumetric projections of confocal images taken from the DGCs area (Supplementary Fig. 6a). The volume of WFA-stained region was greatly increased in the TMEV group (Supplementary Fig. 6b), however, the mean puncta of vGLUT1 was unchanged compared to the sham-injected control mice (Supplementary Fig. 6c). Similarly, in another set of immunostaining, the mean puncta of vGAT in the DGCs from TMEV-infected mice was comparable to the control group (Supplementary Fig. 6d-f). These results show no change in the presynaptic excitatory and inhibitory innervation of DGCs during TMEV-induced acute seizure period and rule out ECM-mediated synaptic reorganization as a contributing factor for increased frequency of postsynaptic currents recorded from mice with seizures. Alternatively, increased mIPSC frequency might have resulted from an increased probability of neurotransmitter release after TMEV infection, however, this may not be the case in the absence of an increase in sIPSC frequency and amplitude. In contrast, the frequencies of both mEPSC and sEPSC were increased in mice with seizures, which could be a consequence of an increased expression of CSPGs in the DG because the negatively charged CSPGs are known to attract calcium ions^41^. Consequently, that could enhance extracellular concentration of Ca^2+^ ([Ca^2+^]), increase presynaptic availability of Ca^2+^, which in turn, may increase the release of glutamate and cause hyperexcitation.

### Negatively charged CSPGs retain K^+^ explaining the observed membrane depolarization

TMEV-infected WT C57BL6/J and Acan^+/+^ mice are highly susceptible to seizures and present with enhanced deposition of CSPGs in the DG. The electrophysiological studies revealed that the DGCs from both WT C57BL6/J and Acan^+/+^ mice with seizures are hyperexcitable compared to non-infected control and TMEV-infected DG-AMG-Acan^−/−^ mice, respectively. Furthermore, the results above also show that an enhanced deposition of CSPGs in the DG and amygdala directly contributes to seizures. This raised questions whether an enhanced deposition of CSPGs in the DG augments seizures by causing changes in the RMP, firing rate, and synaptic currents recorded from the DGCs.

The RMP of DGCs from mice with seizures was significantly depolarized compared to the control mice, which could contribute to high firing rate of these cells. RMP is strongly influenced by the concentration gradients of potassium and sodium ions across the neuronal membrane. Given the highly negatively charged nature of CSPG molecules, they have been speculated to function as an ion exchanger with a strong buffering tendency for cations^14, 15^. Increase in extracellular concentration of K^+^ ([K^+^]⍰) is known to depolarize RMP of neurons and cause hyperexcitability^42^. Furthermore, [K^+^]⍰ is often elevated during seizures in animals and patients with epilepsy^42^. Therefore, we questioned whether enhanced expression of CSPGs around DGCs could depolarize their RMP by locally increasing [K^+^]⍰.

Since the native ECS in hippocampal brain slices is too small to be accessed by ion selective probes without damage to the neurons and their surrounding CSPGs, we instead addressed this question by simulating the interaction of CSPGs with K^+^ in a dish using chondroitin sulfate (CS) agarose gel prepared in 3 mM KCl solution and by measuring [K^+^] in the gel using a K^+^-sensitive microelectrode (Supplementary Fig. 7a). The voltage recorded by the K^+^-sensitive microelectrode is proportional to the available [K^+^] that can interact with the potassium ionophore at the tip of the electrode and the absolute change in [K^+^] can be derived from the calibration curve for each electrode^43^. The experiment posits that CS, by electrostatic interaction with K^+^ ions in solution, will reduce the availability of free K^+^, and that increasing concentrations of CS will bind up increasing K^+^ hence resulting in a lower availability of K^+^ to be sensed by the electrode. CS agarose gels were prepared in 3 mM KCl solution to simulate physiological K^+^ and contained different concentrations of CS. Representative changes in voltage recorded after repeatedly inserting the electrode into 0%, 0.5%, and 5% CS agarose gels are shown in Supplementary Fig. 7b. The voltage change recorded from the control gel (0% CS) was close to zero (Supplementary Fig. 7c). In contrast, significant negative shifts in voltage were recorded in gels with 0.5% and 5% CS. [K^+^] was calculated from the voltage change recorded using the Nernst equation, which showed significant reduction in 0.5% and 5% CS gels (Supplementary Fig. 7d, e). The relative change in [K^+^] suggests that 5% CS in gel is able to change free [K^+^] by 1 mM, from 3.14 to 2.14 (Supplementary Fig. 7e). As this decrease reflects the “retention” of K^+^ to the gel, in the vicinity to the cell, it would equate to a relative 1 mM increase in stationary K^+^ on the outer cell surface. A 1 mM increase in [K^+^]⍰ would result in 7.3 mV depolarization of the membrane potential, which is close to that recorded from DGCs (6.8 mV) of TMEV S+ mice (Fig. 4b; Sham -71.8 mV, TMEV S+ -65 mV). It is concluded from these data that CS can indeed capture K^+^ and increases overall [K^+^]⍰ explaining the observed neuronal depolarization.

## Discussion

Studies of human epileptic brain and animal models of epilepsy suggest that seizures can change the structure and composition of ECM^16^, and on the contrary, altering ECM can also lead to seizures^44^, yet the mechanistic contributions of ECM changes to epilepsy remains elusive. There is emerging evidence that compact PNNs that enwrap soma and proximal processes of GABAergic interneurons regulate excitation-inhibition balance by supporting the inhibitory functions of interneurons^45^. PNNs promote maturation of PV+ interneurons, facilitate their high firing rate by decreasing their membrane capacitance and potentially by regulating ionic reversal potentials and ion channel conductances^13, 46^. Motivated by findings in tumor associated epilepsy, where MMPs-mediated proteolysis of PNNs explained the cortical hyperexcitability by a change in the firing rate of PV+ interneurons^13^, the current study examined viral infection-induced epilepsy where the major change in ECM was the *de novo* appearance of CSPGs filling the interstitial spaces around excitatory neurons in the DG and amygdala. Increases in ECM components have been reported after CNS insults-induced tissue damage^47, 48^, yet the morphology of DG and amygdala was not adversely affected by TMEV infection. Interestingly, we also found increased expression of CSPGs in the DG and amygdala in the pilocarpine-induced SE model of epilepsy. Moreover, similar enhanced deposition of CSPGs in the morphologically intact DG have also been reported in a genetic mouse model of Angelman syndrome that is highly susceptible to flurothyl kindling-induced chronic seizures^49^, in kainic acid-induced SE model of epilepsy^50^, and in the genetic models of epilepsy caused by deletion of bassoon^50^ and Kv1.1^51^. Hence the common upregulation of CSPGs in the DG across these various models of epilepsy are highly suggestive of a common underlying contribution of CSPGs to seizure etiology. While the physiological consequences of increased CSPGs in the DG were not investigated in these genetic epilepsy models, our study clearly suggests a causal role for CSPGs in seizure etiology. Notably, preventing upregulation of CSPGs in the DG and amygdala using Acan^fl/fl^ mice shows a reduction in seizure burden upon aggrecan deletion. Moreover, our electrophysiological data show that the DGCs from WT mice with seizures are hyperexcitable as they exhibit depolarized RMP, high firing rate, and increased synaptic currents, and this hyperexcitability can be rescued by inhibiting *de novo* synthesis of CSPGs in the DG and amygdala.

How might enhanced CSPGs in the DG cause DGCs to be hyperexcitable? It has been known for over sixty years that CS binds to biologically relevant cations (K^+^, Na^+^, Ca^2+^) due to its high anionic charges^52^. The density of fixed anionic charges in the ECM surrounding neurons has been measured in the range of 0.4 to 0.5 M^14^. The presence of such high fixed anionic charges creates a Gibbs-Donnon condition driving electrostatic interactions with ions affecting their transmembrane distribution. We reason, based on this argument, that enhanced deposition of CSPGs in the DG could create a stationary layer of increased extracellular K^+^ which would depolarize RMP, and increase synaptic neurotransmission by accumulating cations in the ECS immediately around DGCs. It can be imagined that the binding of extracellular K^+^ to CSPGs creates electrochemical gradient for K^+^ that draws K^+^ outward from the neurons through K^+^ leak channels until equilibrium sets and depolarizes reversal potential of K^+^. Since constitutively activated K^+^ channels are the prime factor setting up the RMP, increased K^+^ accumulation in the ECS significantly depolarizes the RMP^42^. Our results from K^+^ recording in CS gels support this argument. Furthermore, as per the surface potential or the Guoy-Chapman-Stern theory, the extracellular cations screen negative charges of phospholipids on the outer layer of the neuronal membrane, form diffuse electric double layer and partly reverse the attenuation in membrane potential caused by the membrane-associated negative charges^53, 54^. The binding of extracellular K^+^ by CSPGs could unscreen these phospholipid negative charges, and thus, could also contribute to depolarizing the RMP. CSPGs can also similarly increase extracellular Na^+^, however, the extent of such increase would not be sufficient to achieve supraphysiological level of extracellular Na^+^ required to have any significant impact on the RMP.

Since CSPGs are major constituent of PNNs as well, it would be expected that the PNNs depolarize the RMP of interneurons that they enwrap. However, previous studies have not found changes in the RMP of fast-spiking interneurons upon pathological or experimental degradation of PNN^13, 55^. The effects of ECM degradation are not uniform across different brain regions and the types of neurons affected. For example, enzymatic degradation of PNNs in the brain slices from mice causes plasticity of excitatory synapses in area CA2^56^, whereas suppresses in area CA1^57^. We had found that PNNs decrease membrane capacitance of fast-spiking PV+ interneurons in the cortex^13^. In contrast, the present study shows no change in the membrane capacitance of DGCs upon changes in the CSPG level in the DG. The reasons for these discrepancies are not clearly understood, however, the structure of ECM and the type of neuron appear to play a determining role. We propose a unifying explanation in Fig. 7 that describes how structural differences in perineuronal ECM could differentially influence neuronal physiology. Specifically, the perineuronal ECM around inhibitory interneurons forms a classic PNN structure (Fig. 7b). Due to its high density and being juxtaposed to the neuronal membrane, the structure largely retards ion diffusion within the PNN but rather serves as an insulator that increases the thickness of the membrane and thereby increases charge separation across the membrane (Fig. 7a, left). Consequently, it reduces the membrane capacitance allowing cells to fire at higher firing rates. This arrangement is akin to myelin sheath where a decrease in capacitance plays a major role in accelerating conductance along the myelinated axons. By contrast, the perineuronal ECM around excitatory neurons, such as DGCs, is less compact than the PNNs surrounding interneurons (Fig. 7c) and readily permits ions diffusion within and binding to negatively charged residues on the ECM. Due to the permissibility of ion diffusion within, this ECM does not increase the charge separation across the neuronal membrane. However, it attracts cations and increases their local concentration in the extracellular compartment near the neuronal membrane (Fig. 7a, right). Increased [K^+^] in turn depolarizes the membrane causing these excitatory neurons to be hyperexcitable.

**Fig. 7.**
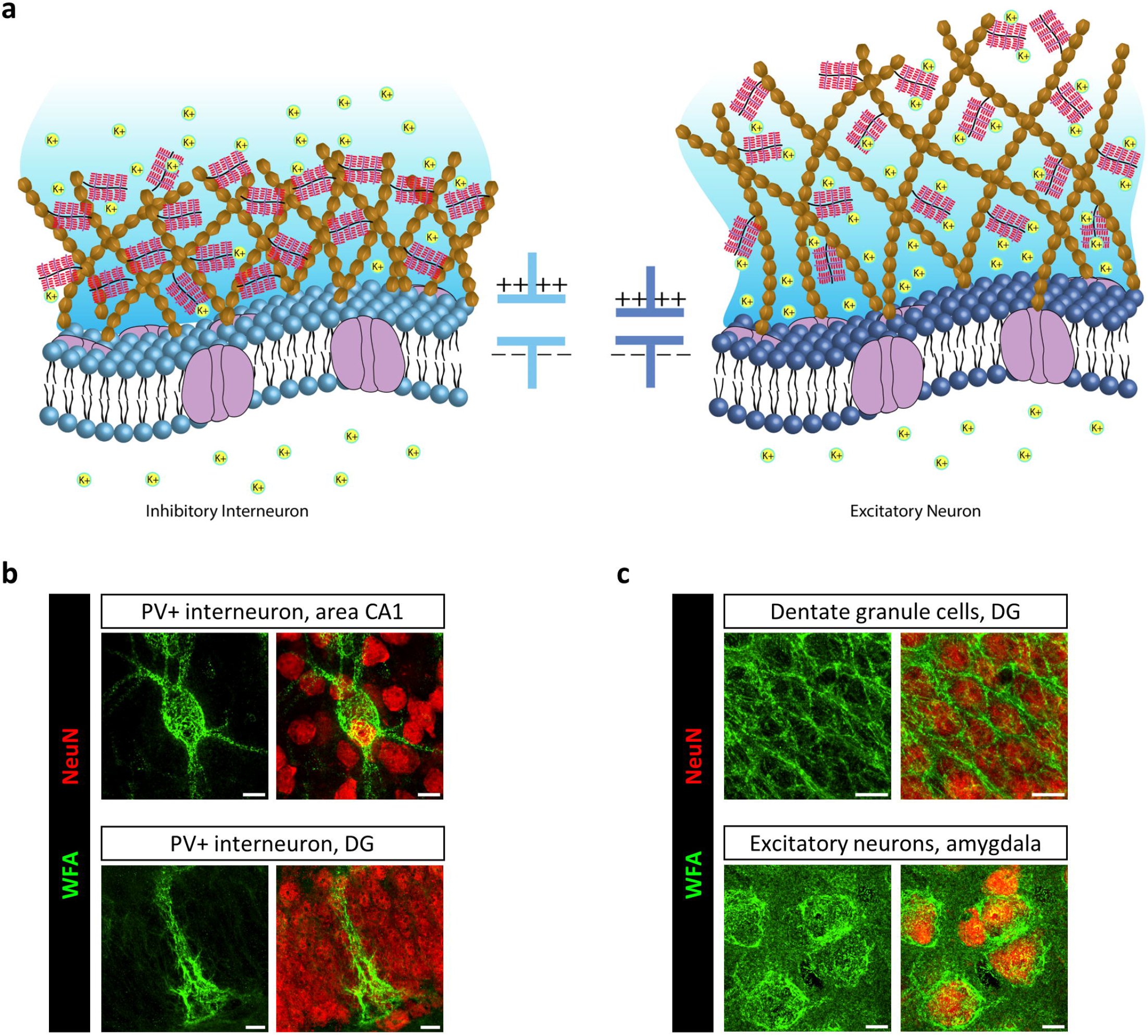
Structural heterogeneity of perineuronal ECM differentially influences neuronal physiology. **a** Left: Perineuronal ECM around inhibitory interneurons forms a condensed tightly woven lattice-like structure enwrapping soma as well as proximal dendrites. This arrangement of dense ECM in close proximity to the neuronal membrane retards ion diffusion and functions akin to dielectric material or myelin sheath and increases the charge separation between intracellular and extracellular compartments. As a result, it decreases membrane capacitance and allows interneurons to fire at higher frequency. Right: Perineuronal ECM around excitatory neurons is less compact and mostly present around soma. The lower density of ECM allows ionic diffusion and does not increase charge separation across the membrane, and therefore, does not affect the membrane capacitance. Instead, it acts akin to sponge attracting cations, such as K^+^, and raises their concentration in the extracellular compartment near the membrane. Increased [K^+^] consequently causes hyperexcitability by depolarizing the resting membrane potential of excitatory neurons. **b** Examples of perineuronal ECM surrounding parvalbumin-containing (PV+) inhibitory interneurons from area CA1 and DG from TMEV/sham-injected mice showing a condensed PNN surrounding soma and proximal dendrites. **c** Examples of perineuronal ECM around excitatory neurons from the DG and amygdala from TMEV-infected mice. The structure of ECM is distinctly different than PNNs around PV+ interneurons. The images shown in panels b and c are maximum intensity projection of z-stack confocal images from the brain slices stained with markers for CSPG (WFA) and neurons (NeuN). Scale bar – 10 µm.

Ionic interactions and binding of CSs with divalent cations, e.g., Ca^2+^, is even greater than for monovalent cations^41, 53^. CSs have been shown to shift the activation curve of unspecified voltage-gated calcium channels towards lower voltage and to increase calcium currents in *Xenopus laevis* photoreceptors, as expected from the Guoy-Chapman-Stern theory and the calcium binding properties of CSs^54^. Furthermore, synaptic release of neurotransmitters is highly sensitive to [Ca^2+^] ^58^. Increase in the [Ca^2+^] causes an increase in the presynaptic basal [Ca^2+^], which in turn, increases the occupancy of synaptic vesicle docking sites and enhances the release probability of neurotransmitters^59^. Thus, CSPGs-mediated enhancement of [Ca^2+^] could increase the availability of presynaptic Ca^2+^ and thereby facilitate neurotransmitter release. However, experimental evidence is needed to substantiate this mechanism contributing to TMEV-induced acute seizures.

Although CSPGs-mediated ionic regulation explains the results of the present study, we do not rule out alternative mechanisms. Given that CSPGs can directly alter the functions of ion channels and the complexity of its effects on neuronal physiology, it is possible that the changes in the expression and/or activities of ion channels as a direct result from enhanced CSPGs could have partly contributed to the results. Further studies are warranted to understand the dynamics between the ECM changes and ion channels functions during seizure development.

In conclusion, the present study reveals increased expression of CSPGs in the DG and amygdala as one of the causal factors for TMEV-induced epilepsy. The excessive CSPGs surrounding the excitatory cells renders these cells hyperexcitable by increasing the concentrations of cations, especially K^+^ and Ca^2+^, in the ECS around these neurons which depolarizes their RMP and increases net excitatory synaptic currents from these cells. Similar enhancements of CSPGs in the DG have been reported in the chemoconvulsant-induced acquired epilepsy models as well as in the genetic epilepsy models suggesting that the enhancement of CSPGs in the DG as a common histopathological phenomenon causing seizures.

## Methods

### Animals (procurement, housing, breeding, and ethics)

C57BL/6J (strain # 000664), FVB/NJ (strain # 001800), B6.FVB(Cg)-*Mmp9*^tm1Tvu^/J (MMP9^−/−^mice, strain # 007084), and B6.FVB-Tg(Pomc-cre)1LowI/J (strain # 010714; hereafter referred to as POMC-cre mice) were purchased from the Jackson laboratories, USA. C57BL/6N-Acan^tm1c^(EUCOMM)^Hmgu^/H (European Mouse Mutant Archive stock # EM:10224; hereafter referred to as Acan^fl/fl^ mice), which contains targeted insertion of loxP sites flanking exon 4 of *acan* gene, were purchased from the Mary Lyon Centre at Medical Research Council Harwell, UK. Mice were housed in groups with a maximum of five mice per cage in an environmentally controlled vivarium providing 12 hr of light/dark cycles starting at 6:00 AM. Mice had free access to food and water *ad libitum*. Mice were allowed to acclimatize for at least 3 days upon arrival at our facility before using them for the experiments. POMC promoter drives expression of cre recombinase strongly in the DGCs^35^. A three-staged breeding strategy was employed to achieve targeted deletion of *acan* gene in the DGCs. First, male hemizygous POMC-cre mice were mated with female homozygous Acan^fl/fl^ mice to get offspring heterozygous for loxP and either hemizygous or noncarrier for cre. Second, mice heterozygous for loxP and hemizygous for cre were backcrossed with homozygous Acan^fl/fl^ mice to get about 25% of the progeny Acan^fl/fl^/POMC-cre (experimental group – homozygous for loxP and hemizygous for cre) and about 25% of the progeny Acan^fl/fl^/noncarrier (control group – homozygous for loxP and noncarrier for cre). Lastly, Acan^fl/fl^/POMC-cre mice were mated with Acan^fl/fl^/noncarrier mice to maintain a single colony containing about half of Acan^fl/fl^/POMC-cre mice and the remaining Acan^fl/fl^/noncarrier mice. All the mice were genotyping as per the PCR protocol provided by the Jackson laboratories and the Medical Research Council Harwell. Both male and female mice were included in all the experiments conducted. All the procedures performed were in accordance with the National Institutes of Health (NIH) Guide for the Care and Use of Laboratory Animals and approved by the Institutional Animal Care and Use Committee of the University of Virginia.

### Procedure of TMEV infection and handling-induced seizure monitoring

Daniel’s strain of TMEV was used to induce seizures in mice. TMEV was kindly provided by the laboratories of Drs. Karen S. Wilcox and Robert S. Fujinami from the University of Utah. The titer of the stock used was 2x10^7^ plaque forming units (PFU) per ml. Mice were anesthetized using 2-3% isoflurane during the entire infection procedure. The head surface over the right hemisphere was disinfected with alcohol swab and the injection site was identified slightly medial to the equidistant point on the imaginary line connecting the right eye and the right ear. Mice were injected with 20 µl of either phosphate-buffered saline (PBS) or TMEV solution (2x10^5^ PFU) intracortically by inserting a 28-guage needle perpendicular to the skull surface. A sterilized syringe containing a plastic sleeve on the needle to expose only 2.5 mm of needle from the tip was used to restrict the injection in a cortical region about 2-3 mm lateral and 1-2 mm posterior to bregma without damaging the hippocampus. The needle was kept in place undisturbed for about 1 min after injection and retracted slowly to minimize leakage. The injury site was disinfected post-injection. Mice resumed their normal behavior within 5-10 min of the procedure.

TMEV-infected mice experience handling-induced acute behavioral seizures between 2-8 dpi^36^. Mice were briefly agitated by shaking their cage and observed for behavioral seizures for about 5 min in two observation sessions daily separately by at least 2 hr. Most of the acute seizures occurred within a minute of handling the mice. Seizure severity was graded using a modified Racine scale as follows: stage 1, mouth and facial movements; stage 2, head nodding; stage 3, forelimb clonus; stage 4, forelimb clonus, rearing; stage 5, forelimb clonus, rearing, and falling; and stage 6, intense running, jumping, repeated falling, and severe clonus^60^. For some studies, seizures were also monitored by video-electroencephalography (vEEG).

### Pilocarpine-induced status epilepticus model of epilepsy

All the required chemicals for this procedure were obtained from MilliporeSigma. FVB/NJ mice (7-8 weeks old) were treated with either pilocarpine (300 mg/kg, intraperitoneal (i.p.)) or saline (0.9% sodium chloride solution) 30 min after methylatropine bromide (5 mg/kg, i.p.) treatment. Mice were video-monitored continuously. Most of the pilocarpine-treated mice developed SE within 5-10 min of pilocarpine injection. SE was identified as a period of continuous behavioral hyperexcitability lasting for over half an hour. Mice with SE exhibited several limbic seizures (stage 3 or greater on the Racine scale) during the period of persistent head nodding. Diazepam (10 mg/kg, i.p.) was administered 2 hr after the onset of stage 3 or greater seizures, and repeated if convulsions were not resolved within 2 hr of the first injection. If convulsions were not resolved within one hour of the second diazepam injection, mice were euthanized. Mice were injected lactated Ringer’s solution subcutaneously and kept warm with a heating pad to facilitate recovery. Mice were monitored for body weight, seizure, and overall health daily until they were sacrificed at 5 days post-SE for the experiment.

### Minocycline treatment

For pharmacological inhibition of MMPs, mice were treated with minocycline (50 mg/kg at 10 ml/kg) intraperitoneally every 12 hr for 8 days starting 2 hr before TMEV/sham infection. Minocycline hydrochloride (#M9511) was obtained from Millipore Sigma. A stock solution of minocycline in saline (5 mg/ml) was prepared fresh just before use. Since minocycline is not readily soluble in saline at 5 mg/ml concentration, the solution was rigorously vortexed, sonicated for 5-10 min and kept in waterbath at 32-37 °C until fully dissolved. The pH and osmolality of solution were set at the physiological level before use. Mice felt discomfort at the injection site for a few seconds immediately after injections but recovered quickly.

### Gel electrophoresis and western blot

Protein levels of aggrecan fragments formed by MMP-mediated cleavage of aggrecan core protein were measured by Western blot analysis. TMEV- and sham-treated mice were sacrificed at 5 dpi. The ipsilateral and contralateral hippocampi were rapidly dissected out and collected separately. All the tissue samples were flash-frozen using 2-methylbutane chilled on dry ice and stored at -80 °C until further processing. The hippocampi samples were homogenized using a rotor-stator homogenizer in a lysis buffer (50 mM Tris-HCl pH 8.00, 150 mM NaCl, 1% Triton X-100, 0.5% sodium deoxycholate, 0.1% sodium dodecyl sulfate (SDS), protease inhibitors (P8340, Sigma) and phosphatase inhibitors (P0044, Sigma); 10 µl lysis buffer per mg of tissue), and the supernatant was collected after centrifugation (15,000 g, 20 min, 4 °C). Total protein concentration in supernatant was determined using bicinchoninic acid (BCA) protein assay (Pierce^TM^ BCA Protein Assay Kit, Catalog #23225, ThermoFisher Scientific). 30-40 µg total protein sample was denatured at 75 °C for 7.5 min and then electrophoresed using polyacrylamide gel (4–15% tris-glycine extended polyacrylamide gel, #567-1085, Bio-Rad) under denaturing conditions. The proteins were transferred to a PVDF membrane which was then blocked with tris-buffered saline-based Odyssey blocking buffer (LI-COR) for 1 hr at room temperature (RT) and incubated with mouse monoclonal anti-aggrecan neoepitope antibody (10 µg/ml; clone AF-28, MAB19310-C, Millipore) and rabbit polyclonal anti-α-tubulin antibody (2 µg/ml; 2144, Cell Signaling Technology) overnight at 4 °C. After washing the primary antibodies, the membrane was incubated with secondary antibodies (IRDye® 800CW donkey anti-mouse IgG, 0.2 µg/ml, 925-32212 and IRDye® 680RD donkey anti-rabbit IgG, 0.2 µg/ml, 925-68073; LI-COR) for 2 hr at RT. The membrane was then imaged using Odyssey imaging system (LI-COR) after washing the secondary antibodies. Densitometric analysis of protein levels was performed using Image Studio software (LI-COR).

### Gel zymography

Gel zymography was conducted to measure the enzymatic activity of MMPs using Novex™ 10% Zymogram Plus (Gelatin) protein gels (ZY00105BOX, ThermoFisher Scientific) which contains 10% gelatin. MMP2 and MMP9, which use gelatin as substrate, are detected by this method. 10 µg total protein sample of hippocampal homogenate was denatured in SDS buffer under non-reducing conditions without heating and electrophoresed using Novex™ 10% Zymogram Plus gel under denaturing condition. MMPs in the gel were renatured by incubating gel in renaturing buffer (50 mM Tris-HCL, 2.5% Triton X-100, 5 mM CaCl_2_, 1 µM ZnCl_2_, pH 7.5) for 30 min twice at RT with gentle agitation. Gel was then equilibrated in developing buffer (50 mM Tris-HCL, 1% Triton X-100, 5 mM CaCl_2_, 1 µM ZnCl_2_, pH 7.5) for 5-10 min at 37 °C and incubated in fresh developing buffer at 37 °C for 48 hr. Next, the developing buffer was discarded, and the gel was stained with a staining solution (aqueous solution containing 0.5% w/v Coomassie brilliant blue G-250, 40% v/v methanol and 10% v/v acetic acid) for 1 hr at RT, washed with deionized water, and destained using a destaining solution (aqueous solution containing 40% v/v methanol and 10% v/v acetic acid) for 30 min at RT with gentle agitation. The destaining step was repeated as necessary to visualize clear bands (caused by digestion of gelatin by MMPs) against a dark blue background. The gel was then imaged using ChemiDoc MP Imaging System (Bio-Rad). Densitometric analysis of protein levels was performed using Image Studio software (LI-COR).

### In situ zymography

TMEV/sham-treated mice were sacrificed at 5 dpi to collect brains for in situ zymography. Mice were deeply anesthetized by i.p. injection (10 ml/kg of body weight) of an aqueous solution of 200 mg/kg ketamine and 20 mg/kg xylazine in saline. After confirming no pedal reflex in response to firm toe pinch, mice were exsanguinated by transcardially perfusing with ice-cold PBS for 2 min. The brain was quickly dissected out and collected in ice-cold N-methyl-D-glucamine (NMDG)-based protective artificial cerebrospinal fluid (ACSF) saturated with 95% oxygen and 5% carbon dioxide to preserve the enzymatic activity. To achieve rapid freezing of tissue without forming ice crystals, the brain was dissected coronally into rostral and caudal parts and the cerebellum was trimmed out to reduce the tissue size. Brain tissues were cryopreserved by immersing them in tissue freezing medium (TFM, Electron Microscopy Sciences) in a plastic mold (4566, Tissue-Tek Cryomold). The molds were then rapidly frozen by submerging in a beaker filled with chilled 2-methylbutane on dry ice. The entire process of placing the tissue into TFM and snap-freezing it was completed within 1.5 min, which ensured the preservation of enzymatic activity. The frozen tissue blocks were stored at -80 °C and moved to -20 °C a day before cryosectioning. 15 µm thick coronal slices were prepared using cryostat (CM1850 UV Cryostat, Leica) at -20 °C and mounted on glass slides (48311-703, VWR Superfrost Plus Micro Slide). The glass slides were kept on dry ice and stored at -80 °C until further processing.

Proteolytic activity of MMPs was measured in brain slices using EnzChek Gelatinase/Collagenase Assay Kit (E-12055, ThermoFisher Scientific). This kit contains fluorogenic DQ gelatin (highly quenched fluorescein-labeled gelatin) as a substrate mainly for MMP2 and MMP9. The enzymatic digestion of this substrate yields green fluorescence proportional to the endogenous activity of MMP2/9. Briefly, the slices were thawed on ice for 30 min and washed 3-4 times with PBS to remove TFM. The slices were then reacted with a solution of DQ gelatin fluorescein conjugate (20 µg/ml) at 37 °C for 1 hr in a dark and humid chamber for optimal enzymatic activity. Next the slices were washed 3-4 times with PBS to remove excess fluorogenic substrate and fixed with 4% PFA for 5 min. A mounting media (P36930, ProLong Gold Antifade Mountant, ThermoFisher Scientific) was applied to the slices and a glass coverslip (12-548-5E, Fisher Scientific) was placed over the slices. The edges of the coverslip were sealed with nail polish. The slices were imaged using Nikon A1 confocal microscope and the images were analyzed by NIS-Elements software (Nikon).

### Immunohistochemistry

Mice were anesthetized by injecting (i.p.) a mixture of ketamine and xylazine (200 mg/kg and 20 mg/kg, respectively, 10 ml/kg) and exsanguinated by transcardially perfusing with ice-cold PBS for 2 min followed by 4% PFA for 5 min. The brain was dissected out and further fixed in 4% PFA overnight at 4 °C before storing it in PBS containing 0.02% sodium azide at 4 °C. Next, 50 µm thick coronal slices were prepared using vibratome (5100mz, Campden Instruments). The slices were stored in PBS containing 0.02% sodium azide at 4 °C for short term (<6 months) use, and for a longer-term storage, transferred into cryoprotective medium (118 mM NaH_2_PO_4_, 94 mM NaOH, 2.5 mM HCL, 30% v/v glycerol, 30% v/v ethylene glycol in deionized water, pH 7.2-7.4) and stored at -20 °C.

We batch-processed slices from all mice (two slices from each mouse) in the experimental cohort simultaneously to minimize procedural variability. The slices were washed with PBS 2-3 times, and permeabilized and blocked by incubating in blocking buffer (0.03% v/v Triton X-100 and 10% v/v normal goat serum in PBS) for 1 hr at RT. The slices were then incubated in primary antibodies solution prepared in diluted blocking buffer (1:3 of blocking buffer and PBS) overnight at 4 °C. Next day the slices were washed with PBS 4 times (5 min each), the slices were incubated in secondary antibodies solution prepared in diluted blocking buffer for 2 hr at RT in dark. The slices were again washed with PBS 4 times (5 min each) to remove unbound secondary antibodies. The slices were then mounted on the glass slides (48311-703, VWR Superfrost Plus Micro Slide), allowed to air-dry for about 15-20 min, and covered with a mounting media (P36930, ProLong Gold Antifade Mountant, ThermoFisher Scientific) and glass coverslip (12-548-5P/5M/B, Fisher Scientific). The edges of the coverslip were sealed with nail polish.

Primary antibodies and markers used were biotinylated WFA (4 µg/ml, B-1355-2, Vector Laboratories), guinea pig polyclonal anti-NeuN antibody (1-2 µg/ml; 266004, Synaptic Systems), rabbit polyclonal anti-NeuN antibody (1 µg/ml; ABN78, MilliporeSigma), mouse monoclonal anti-PV antibody (1 µg/ml; PV235, Swant), rabbit polyclonal anti-PV antibody (1 µg/ml; PV27, Swant), guinea pig polyclonal anti-vGLUT1 antibody (2 µg/ml; 135304, Synaptic Systems), and mouse monoclonal anti-vGAT antibody (2 µg/ml; 131011, Synaptic Systems). Alexa Fluor 555-conjugated streptavidin (S32355, ThermoFisher Scientific; 4 µg/ml) was used to detect biotinylated WFA. Appropriate goat secondary antibodies (1-2 µg/ml) from ThermoFisher Scientific and Jackson ImmunoResearch were used to immunodetect primary antibodies.

### Confocal image acquisition and analysis

Images were acquired at different magnifications using Nikon A1 or Olympus RS3000 confocal microscope and analyzed using NIS-Elements AR analysis program (Nikon), Imaris (Oxford Instruments), and Image J (NIH). Imaging parameters such as laser intensity, exposure time, confocal aperture diameter, resolution, step size for the z-stacks, gain and offset values of the detector were set carefully and consistently across the comparison groups for each experimental cohort.

Mean fluorescence intensities of WFA staining in the granular and molecular layers of DG and in amygdala were quantified by drawing region of interests surrounding these brain areas and subtracting mean background fluorescence. At least two brain slices from 4-5 mice each were quantified, and the numbers of brain slices analyzed were considered as n for the statistical tests. Structual integrity analysis of PNN+ neurons was performed by taking fluorescene intensity measurement along a line drawn over PNN as described in detail previously^13^.

Quantification of the volumetric images was performed using Imaris. Volume of WFA and puncta of vGLUT1 and vGAT were calculated using the surface creation tool with smoothing detail enabled and a surface grain size set to 0.103 µm. Background subtraction was enabled with diameter of largest sphere set to 0.388 µm and manual thresholding was set to a value of 200. Noise was removed using a filter surface voxel setting with a lower limit of 10 (for WFA) or 1 (for vGLUT1 and vGAT). Sum volume for WFA was automatically calculated and then recorded from the detailed statistics panel. The number of surfaces for vGLUT1 and vGAT was automatically calculated and then recorded from the overall statistics panel.

### Protective recovery method for acute brain slice preparation

All mice used for patch-clamp electrophysiology were between 4-10 days post-TMEV/sham treatment. Acute brain slices were prepared using an optimized NMDG protective recovery method^61^. This method significantly enhanced brain slice viability and provided excellent preservation of neuronal morphology. All the solutions used in this method were continuously oxygenated with a mixture of 95% oxygen and 5% carbon dioxide. Mice were first anesthetized by injecting a mixture of ketamine and xylazine (200 mg/kg and 20 mg/kg, respectively, 10 ml/kg, i.p.) and exsanguinated by transcardially perfusing with a cold NMDG-based ACSF (concentrations in mM: 92 NMDG, 2.5 KCl, 1.2 NaH_2_PO_4_, 30 NaHCO_3_, 20 HEPES, 25 D-glucose, 5 sodium-L-ascorbate, 3 sodium pyruvate, 0.5 CaCl_2_, and 10 MgCl_2_; pH 7.3-7.4, osmolality 310-320 mOsm/kg) for 1.5 min. The brain was quickly isolated and collected in ice-cold NMDG ACSF to allow it to uniformly cool for about 1 min. The cerebellum and a quarter of the cerebrum from the rostral end were discarded and the remaining brain was sliced horizontally (300 µm thick) using vibratome (VT1200S, Leica) in a cold NMDG ACSF. All the slices were transferred in a quick succession to a beaker containing 100 ml NMDG ACSF maintained at 32 °C in a waterbath and the slices were allowed to recover from the slicing-induced surface damage for 25 min. During this recovery, the concentration of Na^+^ was increased stepwise from 40 mM in NMDG ACSF to 45, 55, 65, 85, and 125 mM by adding 0.25, 0.5, 0.5, 1, and 2 ml of 2 M NaCl solution, respectively, every 5 min starting immediately after initiating the recovery period. The osmolality of NMDF ACSF was increased to 460-470 mM at the end of recovery, however, it did not affect the slice health as reported earlier^61^. The slices were then transferred to a beaker containing HEPES holding ACSF (concentrations in mM: 92 NaCl, 2.5 KCl, 1.2 NaH_2_PO_4_, 30 NaHCO_3_, 20 HEPES, 25 D-glucose, 5 sodium-L-ascorbate, 3 sodium pyruvate, 2 CaCl_2_, and 2 MgCl_2_; pH 7.3–7.4, osmolality 310–320 mOsm/kg) at RT and kept there until used for patch-clamp recordings.

### Patch-clamp electrophysiology

Acute brain slices were cut into half from the midline and individual hemislices were used for the recording. The slice was placed in the recording chamber of electrophysiology rig continuously superfused (2 ml/min) with warmed (32-33 °C) carbogenated (95% oxygen and 5% carbon dioxide) extracellular recording ACSF (concentrations in mM: 125 NaCl, 3 KCl, 1.25 NaH_2_PO_4_, 25 NaHCO_3_, 5 HEPES, 12.5 D-glucose, 2 CaCl_2_, and 2 MgCl_2_; pH 7.3–7.4; osmolality, 310–320 mOsm/kg). Dentate granule cell layer was identified using a microscope (Zeiss Examiner.D1) fitted with 40X water immersion lens and camera (Zeiss AxioCam MRm, AxioVision software). The recordings were obtained from the granule cells in the outer half of the cell layer in the suprapyramidal blade of DG.

Patch pipettes with 4.5-6.5 MΩ tip resistance were prepared from borosilicate glass capillaries (TW150F-4, WPI) using micropipette pullers (HEKA PIP 6 and MDI PMP102). Patch pipettes were filled with intracellular solution of different chemical composition depending on the purpose of the experiment. For all recordings, fluorescent dye Lucifer yellow (25573, Cayman Chemical) was added to intracellular solution (0.4 mg/ml) on the day of experiment to label the patched cells for morphological verification later. Patch pipettes were guided to the target cells using micromanipulator (MP-225, Sutter Instrument). Whole-cell voltage-clamp and current-clamp recordings were amplified and filtered (1 kHz lowpass bassel filter) using Axopatch 200B amplifier (Molecular Devices). Data were acquired using Digidata 1440A digitizer (Molecular Devices) at 2-10 kHz sampling rate and Clampex 10.7 software (Molecular Devices). For voltage-clamp recordings, the membrane potential was clamped at −70 mV. RMP was measured by setting “I = 0 mode” on amplifier immediately after achieving stable whole-cell patch configuration. Cell membrane capacitance was measured by cancelling the capacitive transient current using amplifier after achieving stable whole-cell patch configuration. Input resistance (R_in_) was calculated using the Ohm’s law by measuring a mean change in membrane potential in response – 100 pA current injection (R_in_ = ΔV/-100). The slices were fixed in 4% PFA overnight after the recordings and later stained with biotinylated WFA (8 µg/ml) to image the level of CSPGs in DG and the morphology of patched DGCs.

#### Action potential recording

The excitability of neurons was assessed by generating input/output curve to measure action potential firing frequency. Intracellular recording solution used for AP recording contained (concentrations in mM) 130 potassium gluconate, 5 KCl, 10 HEPES, 0.5 EGTA, 2 ATP (magnesium salt), 0.2 GTP (sodium salt), and 10 disodium creatine phosphate (pH 7.3 with 1 M KOH, osmolality 295-300 mOsm/kg). Current steps in 20 pA increment (1.1 s duration) were applied from -100 pA and the resulting action potentials were analyzed using Clampfit 11.1 software (Molecular Devices). Spikes with at least 20 mV deflection from the baseline were considered for action potential analysis. Action potential threshold current was measured by injecting current gradually increasing from 0 to 600 pA (300 steps, 2 pA increment per step). The minimum current required to induce action potential consistently was identified as action potential threshold current. Most of the cells fired APs within this current injection range used, however, cells that did not fire within this range were assigned firing threshold of 600 pA.

#### Postsynaptic currents recording

Intracellular solution used for recording postsynaptic currents contained (concentrations in mM) 130 cesium gluconate, 5 CsCl, 10 HEPES, 0.5 EGTA, 2 ATP (magnesium salt), 0.2 GTP (sodium salt), 10 disodium creatine phosphate, and 5 QX-314 chloride (pH 7.3 with 1 M CsOH, osmolality, 295-300 mOsm/kg). The reversal potentials for Na^+^/K^+^ and Cl^−^ were calculated as 0 mV and -70 mV, respectively, at 32 °C given the internal and external recoding solutions used. Therefore, EPSC and IPSC were recorded by clamping the cells at -70 mV and 0 mV, respectively, without requiring inhibitors of glutamatergic and GABAergic currents. For recording mEPSC and mIPSC, 1 µM tetrodotoxin (TTX) was used in the external recording ACSF. The effect of TTX on neural network inhibition was confirmed by observing no AP firing after 10 min of TTX application. Recordings of synaptic currents in some cells were verified by observing a complete elimination of EPSC and IPSC using external recording ACSF containing inhibitors of glutamatergic (20 µM CNQX and 50 µM D-AP5) or GABAergic (10 µM bicuculline) currents, respectively. Series (access) resistance (R_a_), input (membrane) resistance (R_m_), and holding currents (I_h_) were monitored throughout the recording. After whole-cell configuration is achieved, 5-10 min were allowed to stabilize R_a_, R_m_ and I_h_ before initiating recordings. Only recordings that did not exhibit substantial changes (<20%) in R_a_ and R_m_ were used for analysis. Total 3.2 min of recordings from each cell were used for analyzing amplitude, frequency, and kinetics parameters by Mini Analysis 6 software (Synaptosoft).

### K^+^ measurement

[K^+^] was measured in CS agarose gel using K^+^-sensitive microelectrodes which were prepared as described previously^43^. Briefly, electrodes (2-3 MΩ tip resistance) were pulled from cleaned borosilicate glass capillaries (TW150F-4, World Precision Instruments), their tips were silanized using 5% dichlorodimethylsilane at 220 °C for 30 min and stored in a desiccator. The electrode tip was filled to 2-3 mm with K^+^ ionophore solution (IE190, World Precision Instruments) by capillary action. Then the electrode was backfilled with 100 mM KCl reference solution while ensuring no bubbles form at the interface of K^+^ ionophore and reference solutions. Each K^+^-sensitive microelectrode was calibrated with 3 mM KCl and 10 mM KCl solutions before using for measuring [K^+^] in CS gel. CS gel was prepared by dissolving 0.3% agarose and 0-5% CS from bovine trachea (230699, Millipore Sigma) in 3 mM KCl solution in a microwave oven and casting the molten mixture into a beaker. A small uniform piece of the gel was sampled for measuring [K^+^] by inserting a stainless-steel hex nut into the gel. Three hex nuts holding 0%, 0.5% and 5% CS gel were placed in a recording chamber of the electrophysiology rig and the chamber was filled with 3 mM KCl solution. The calibrated K^+^-sensitive microelectrode was lowered into the chamber to take stable voltage measurements first in 3 mM KCl solution (baseline) and then in CS gel under current-clamp recording mode at RT. The process was repeated twice to get three successive measurements from each CS gel to calculate mean change in voltage relative to baseline. The mean change in voltage was applied in the Nernst equation to calculate [K^+^].

### Surgical procedure for vEEG and intracerebral nanoinjections

Mouse was anesthetized using 3% isoflurane, provided analgesia (0.1 mg/kg buprenorphine i.p. and 5 mg/kg carprofen i.p.), and its head was affixed into a stereotaxic instrument (David Kopf Instruments). The hair over the skull area was removed using a hair-removal cream, the surgical area was disinfected using iodine and 70% alcohol, and the skull was exposed. Mouse was continuously anesthetized using nasal tubing supplying 1-2% isoflurane throughout the surgical procedure. For vEEG, a total of five holes (three for anchor screws and two for electrodes) were drilled in the skull carefully without causing bleeding. Two electrodes of a three-channel electrode set up (MS333/8-A, Plastics One) were twisted and implanted in the DG-CA3 region of hippocampus using stereotaxic coordinates of 1.8 mm lateral (ipsilateral to the infection site) and 2.00 mm posterior from bregma, and about 2.5 mm ventral from the skull surface to reach the target area at 1.8 mm ventral from the brain surface. The reference electrode was implanted in the contralateral cortical surface (1.0 mm lateral and 1.0 mm posterior from bregma). Three anchor screws were carefully inserted into the skull without damaging the brain surface – the first one anterior to the bregma ipsilaterally, the second one over the left parietal cortex posterior to the reference electrode, and the third one posterior to the bregma ipsilaterally. The electrodes and screws were secured in position using a dental cement and the skin incision was closed using tissue glue.

For the deletion of *acan* gene in amygdala, cre recombinase was expressed in neurons by injecting AAV9 plasmid construct pENN.AAV9.hSyn.HI.eGFP-Cre.WPRE.SV40 (105540-AAV9, addgene) in amygdala bilaterally. The control group of mice received injections of pAAV9-hSyn-EGFP (50465-AAV9, addgene). The calvarium was carefully leveled after exposing the skull to precisely target injection sites in amygdala stereotactically. The position of mouse head was adjusted to set bregma and lambda in the same horizontal plane and the bilateral insertion sites on the skull surface at the same dorsoventral level relative to bregma. The holes were drilled in the skull at bilateral insertion sites. The coordinates used for amygdala injections were −1.4 AP (anteroposterior) and ±3.2 ML (mediolateral) from bregma, and −4.5 DV (dorsoventral) from skull surface. Injections at these coordinates were tested in a pilot study and found targeting amygdala reliably. The stocks of AAV9 constructs were saved in single-use aliquots at -80 °C. The aliquot was thawed on ice and diluted with saline just before use to inject 1.5 x 10^9^ VG (viral genomes) per injection site. Neuros syringe (Model 1701, 65460-06, Hamilton) and 33-gauge needle (65461-02, Hamilton) were used to inject 150 nl solution at 50 nl/min injection speed (Quintessential Stereotaxic Injector, 53311, Stoelting). The needle was checked for any blockage by pumping a small amount of fluid just before inserting into brain. It was also ensured that the needle penetrated brain straight. The needle was kept undisturbed for 4 min post-injection and slowly retracted over 2.5 min to prevent leakage of any fluid. The holes in skull were filled with bone wax and the skin incision was closed using a tissue glue. All surgically operated mice were treated humanely and provided post-operative care as per the NIH guidelines and the institutional animal care protocol.

### vEEG acquisition and seizure analysis

vEEG was initiated after 10-11 days of recovery from the electrode implant procedure. Mice were connected to EEG100C differential amplifier (BIOPAC Systems) using a lightweight three-channel cable with a three-channel rotating commutator (P1 Technologies). The MP160 data acquisition system and AcqKnowledge 5.0 software (BIOPAC Systems, Inc.) were used to record electroencephalogram. M1065-L network camera (Axis communications) and media Recorder 4.0 software (Noldus Information Technology) were used to record the behavior of each mouse. All the cables and electrical components were sufficiently shielded to minimize electrical noise. Video and EEG recordings were automatically synchronized using Observer XT 14.1 software (Noldus Information Technology). EEG signals were bandpass filtered between 0.5 and 100 Hz, amplified, and digited at a sampling frequency of 500 Hz. Mice had access to food and water conveniently during the entire vEEG recording. A baseline recording was acquired for 1 hr before injecting mice with TMEV as described above, and then the recording was continuously acquired until 7 dpi. The EEG and video recordings were reviewed manually by an experimenter blinded to the genotype of mice. Electrographic seizures were defined as rhythmic spikes or sharp-wave discharges with amplitudes at least two times the average amplitude of baseline, frequency at least 2 Hz, and duration at least 5 s. Seizures were also identified by verifying postictal suppression of the baseline EEG activity which typically occurs after seizure but not accompanied by electrographic artifacts associated with mouse behavior other than seizures. At the end of the experiment, mice were sacrificed to collect either 4% PFA-fixed brains for IHC and electrode placement verification or fresh brain regions for biochemical analysis.

### Data presentation and statistics

Datasets with continuous variables are summarized by plotting mean and the standard error of mean as a scatter plot with bar showing individual datapoints, and the datasets with ordinal variables are represented by frequency distribution unless otherwise stated in the specific figures. Some datasets are represented as box and whisker plots where the horizontal line and plus sign within a box indicate median and mean, respectively, of the dataset. The lower and upper ends of the box indicate the first and third quartiles, respectively. The upper and lower whiskers extend to the highest and lowest values in the dataset, respectively. Individual data points are represented by dots. Parametric statistical tests were used if the data were sufficiently normal distributed and variance within groups was sufficiently similar. Experimental designs with two comparison groups were analyzed by a two-tailed unpaired t-test and the designs with more than two comparison groups were analyzed using either one-way or two-way analysis of variance (ANOVA). Multiple pairwise comparisons were performed by appropriate posttest. Two groups with binomial outcome were analyzed by Fisher’s exact test. Survival (percent seizure free) curves were analyzed by log-rank test. Statistical details of experiments are mentioned in the figure legends. The difference between the two groups was considered statistically significant with a p-value less than 0.05. GraphPad Prism 9 and Microsoft Excel software were used for statistical analysis.

## Acknowledgements

We acknowledge Dr. Jenny Raines and Paul Youmans for their assistance in mice colony management. Figs. 2a and 3a were created with BioRender.com. This work was supported by grants from the National Institutes of Health, USA (RO1-NS036692, RO1-NS082851 and RO1-NS052634).

## Author contributions

D.C.P., B.P.T., and H.S. were involved in conception of the work and design of the experiments. D.C.P., N.S., and C.P. were involved in conducting molecular biology studies, IHC, confocal microscopy and image analysis; D.C.P. conducted in vivo work, acquired and analyzed patch-clamp and vEEG recordings; D.C.P., J.L.B., and M.L.O. contributed to extracellular K^+^ measurement using K^+^-sensitive microelectrodes. L.C. and I.K. contributed to graphic design and image analysis, respectively. D.C.P. and H.S. interpreted the results and drafted the manuscript.

