## Supplementary figure legends for "Infection-induced epilepsy is caused by increased expression of chondroitin sulfate proteoglycans in hippocampus and amygdala"

**Supplementary Fig. 1 |** **Degradation of extracellular matrix in CA3 and CA2 areas of hippocampus in TMEV-infected mice with acute seizures.**

**a-b** Representative maximum intensity projections of micrographs acquired from CA3c (a) and CA3a (b) regions of mice infected with saline (Sham) or TMEV (TMEV S+: mice with acute seizures) at 5 dpi. The brain slices were stained with markers for CSPG (WFA, in green) and neurons (NeuN, in red). The images show significant structural impairment in PNN around CA3 neurons and an extensive loss of CSPGs around dendritic processes.

**c** Images acquired from the CA2 region of hippocampus stained with the markers of PNN (WFA in green), neuron (NeuN in red) and parvalbumin+ interneuron (PV in blue). Areas of the images on left demarcated by white squares are magnified on the right show degradation of PNN in TMEV-infected mice with seizures.

**d** Fluorescence intensity measurement along a line drawn over PNN (black dashed lines over PNN+ cells in panel c) shows multiple peaks and valleys in intensity graph reflecting latticed structure of PNN. The black shaded area laid over the intensity graph indicates background fluorescence intensity. Increase in the distance between two adjacent peaks suggests degradation of PNN around PV+ interneurons in TMEV-infected mice with seizures.

**e** Number of fluorescence intensity peaks measured along a line drawn over PNN shows significant decrease in TMEV-infected mice with seizures compared to sham-treated mice and TMEV-infected mice without acute seizures.

Statistics: One-way ANOVA, Tukey’s multiple comparisons test; n=10-11 brain slices from 4-5 mice per group; ***p*<0.01, *****p*<0.0001. Scale bar displayed in one image applies to all images in that panel.

**Supplementary Fig. 2 |** **Increased activity of MMPs causes degradation of aggrecan in the hippocampus of TMEV-infected mice with acute seizures.**

**a** Image of the gel zymogram stained with Coomassie brilliant blue (colors inverted) shows significantly increased digestion of gelatin embedded in the zymogram by MMP2 and MMP9 in the hippocampal homogenate from TMEV-infected mice with acute seizures.

**b-c** Increased mean optical density of bands in gel zymogram shows increased enzymatic activity of MMP2 and MMP9 in the TMEV S+ groups compared to other two groups at 5 dpi. TMEV-infected mice resistant to acute seizures also had slightly increased enzymatic activity compared to the sham-treated mice (Brown-Forsythe and Welch ANOVA, Dunnett’s T3 multiple comparisons test; n=6-10).

**d** In-situ zymography measures localized enzymatic activity of MMP2 and MMP9 by measuring the intensity of fluorescein released from highly quenched fluorescein-labeled gelatin after its proteolytic digestion. Zoomed images of the CA1 region (highlighted in white dashed rectangles in representative micrographs of hippocampus on left) displayed on the right show increased fluorescence in the TMEV S+ group. The scale bars apply to all comparable images in the panel.

**e** Increased fluorescein-labeled area in hippocampal slices from TMEV-infected mice with seizures at 5 dpi (One-way ANOVA, Tukey’s multiple comparisons test; n=4-7 brain slices from 4-5 mice).

**f** Immunoblot shows significant detection of aggrecan fragments generated by MMP activity using aggrecan neoepitope antibody in the hippocampus from the TMEV S+ group at 5 dpi. α-tubulin is detected to normalize the protein loading amount in electrophoresis.

**g-i** Fluor intensity of aggrecan fragments normalized to corresponding α-tubulin blots shows significant increase in mean fluor intensity of aggrecan fragments of 55 kDa (g), 29 kDa (h), and 25 kDa (i) in the hippocampus from TMEV-infected mice with seizures compared to the other groups (One-way ANOVA or Brown-Forsythe and Welch ANOVA, Tukey’s or Dunnett’s T3 multiple comparisons test; n=6-8; ***p*<0.01, ****p*<0.001, *****p*<0.0001).

**Supplementary Fig. 3 |** **No difference in TMEV-induced acute seizures between WT and MMP9^-/-^ C57BL/6J mice.**

**a** An example of electrographic activity recorded from hippocampus during acute TMEV infection period corresponding to spontaneous behavioral seizure. Segments identified by numbers are magnified below temporally showing the pattern of spikes (1), high frequency discharges during ictal period (2), postictal spiking (3), and postictal suppression (4).

**b** Heatmap shows spontaneous seizures recorded by vEEG based on their severity score for each mouse between 3-6 dpi.

**c** Percentage of total infected mice in each group that remained seizure-free each day. None of the mice developed seizures before 3 dpi.

**d** Average number of seizures per day between 3-7 dpi plotted for each mouse shows no difference in seizure frequency between WT and MMP9^−/−^ mice.

**e** Average cumulative seizure burden, which is calculated as a mean of the summation of all seizure scores for each mouse up to each dpi, shows no difference in seizure severity between WT and MMP9^−/−^ mice (data shown as mean±SEM, n=8).

**f** Mean seizure duration shows no difference between WT and MMP9^−/−^ mice (data shown as mean±SEM, n=6). Each dot represents seizure duration of a single seizure measured from electrographic activity in EEG.

**g** Mean seizure score at each dpi shows no difference between WT and MMP9^−/−^ mice (data shown as mean±SEM, n=8).

**h** Distribution of seizures based on seizure severity score.

**Supplementary Fig. 4 |** **Minocycline significantly inhibits the MMP-mediated degradation of aggrecan in hippocampus in TMEV-infected mice with acute seizures.**

**a** Immunoblot probing for aggrecan fragments generated by MMP activity using aggrecan neoepitope antibody shows significant reduction in the hippocampus from TMEV-infected mice treated with minocycline at 7 dpi. The sham-injected control mice show very low levels of aggrecan fragments. α-tubulin is detected to normalize the protein loading amount in electrophoresis.

**b-d** Fluor intensity of aggrecan fragments normalized to that of α-tubulin blots shows significant decrease in mean fluor intensity of aggrecan fragments of 55 kDa (b), 29 kDa (c), and 25 kDa (d) in the hippocampus from TMEV-infected mice treated with minocycline at 7 dpi (One-way ANOVA, Tukey’s multiple comparisons test; n=5-7; ***p*<0.01, ****p*<0.001, *****p*<0.0001).

**Supplementary Fig. 5 |** **Increased expression of CSPGs in dentate gyrus and amygdala in pilocarpine-treated mice with seizures.**

**a** Comparative images of the coronal brain hemislices obtained from mice treated with saline (Sham) or pilocarpine (PILO) at 5 days post-SE and stained with the markers for CSPG (WFA, in green) and neuron (NeuN, in red).

**b** Enlarged views of hippocampus show a stark increase in the level of CSPGs in DG from the PILO group. Neuronal loss is noticeable in the CA1, CA3a, and CA3b regions from the PILO group.

**c** Enlarged views of amygdala show substantial increase in the level of CSPGs in the PILO group.

**d-f** Image analysis shows a significant increase in mean fluorescence intensity of WFA in the granular and molecular layers of DG (DG-GL (d) and DG-ML (e), respectively) and amygdala (f) from the PILO group compared to the Sham group.

Statistics: Unpaired two-tailed or Welch’s t test; n=10-12 brain slices from 4-5 mice per group; ***p*<0.01, ****p*<0.001, *****p*<0.0001. Scale bar displayed in one image applies to all images in that panel.

**Supplementary Fig. 6 |** **Enhancement of CSPGs in dentate gyrus during acute infection period is not associated with any change in the density of presynaptic vGLUT1 and vGAT puncta.**

**a** Representative volumetric projections of micrographs (17 optical sections, 0.5 µm z-stack step size) acquired from dentate granular cells region of mice infected with saline (Sham) or TMEV (TMEV S+ : mice with seizures) at 5 dpi. The brain slices were stained with markers for CSPG (WFA, in green) and glutamatergic presynaptic boutons (vGLUT1, in red). The scale bar applies to all images in the panel.

**b** Image analysis shows a significant increase in mean volume of WFA fluorescence in the granular layer of DG from TMEV-infected mice (Unpaired two-tailed t test; n=10-11 brain slices from 4-5 mice per group; *****p*<0.0001).

**c** No difference in mean puncta of vGLUT1 between sham- and TMEV-infected mice.

**d** Representative volumetric projections of micrographs (17 optical sections, 0.5 µm z-stack step size) acquired from dentate granular cells region of mice infected with saline (Sham) or TMEV (TMEV S+ : mice with seizures) at 5 dpi. The brain slices were stained with markers for CSPG (WFA, in green) and GABAergic presynaptic boutons (vGAT, in red). The scale bar applies to all images in the panel.

**e** A significant increase in mean volume of WFA fluorescence in the granular layer of DG from TMEV-infected mice (Unpaired two-tailed t test; n=7-8 brain slices from 4-5 mice per group; ***p*<0.01).

**f** No difference in mean puncta of vGAT between sham- and TMEV-infected mice.

Scale bar displayed in one image applies to all images in that panel.

**Supplementary Fig. 7 |** **Negatively charged chondroitin sulfate (CS) binds to K^+^ and reduces the concentration of K^+^ ([K^+^]) in CS agarose gel.**

**a** Experimental set up of a patch-clamp electrophysiology rig to measure K^+^ concentration using K^+^-sensitive microelectrode in CS agarose gel prepared and kept in 3 mM KCl external solution.

**b** Representative current-clamp recordings obtained from CS agarose gels (0-5%). The baseline voltage was recorded from external solution above the CS gels. The deviations in the recordings from the baseline correspond to insertions of K^+^-sensitive microelectrode into the gel.

**c** Mean changes in voltage recorded from CS agarose gels (0-5%) show significant difference between 0% (control), 0.5%, and 5% CS agarose gels.

**d** [K^+^] as calculated from the voltage changes recorded using the Nernst equation show a significant decrease in 5% CS gel compared to control and 0.5% CS gels.

**e** Summary of mean [K^+^] measured in CS agarose gels. (S.D. – standard deviation)

Statistics (c-d): One-way ANOVA, Holm-Šidák’s multiple comparisons test; n=4; **p*<0.05, ****p*<0.001, *****p*<0.0001).
