## Supplementary figures and images for "Infection-induced epilepsy is caused by increased expression of chondroitin sulfate proteoglycans in hippocampus and amygdala"

### Supplementary Fig 1

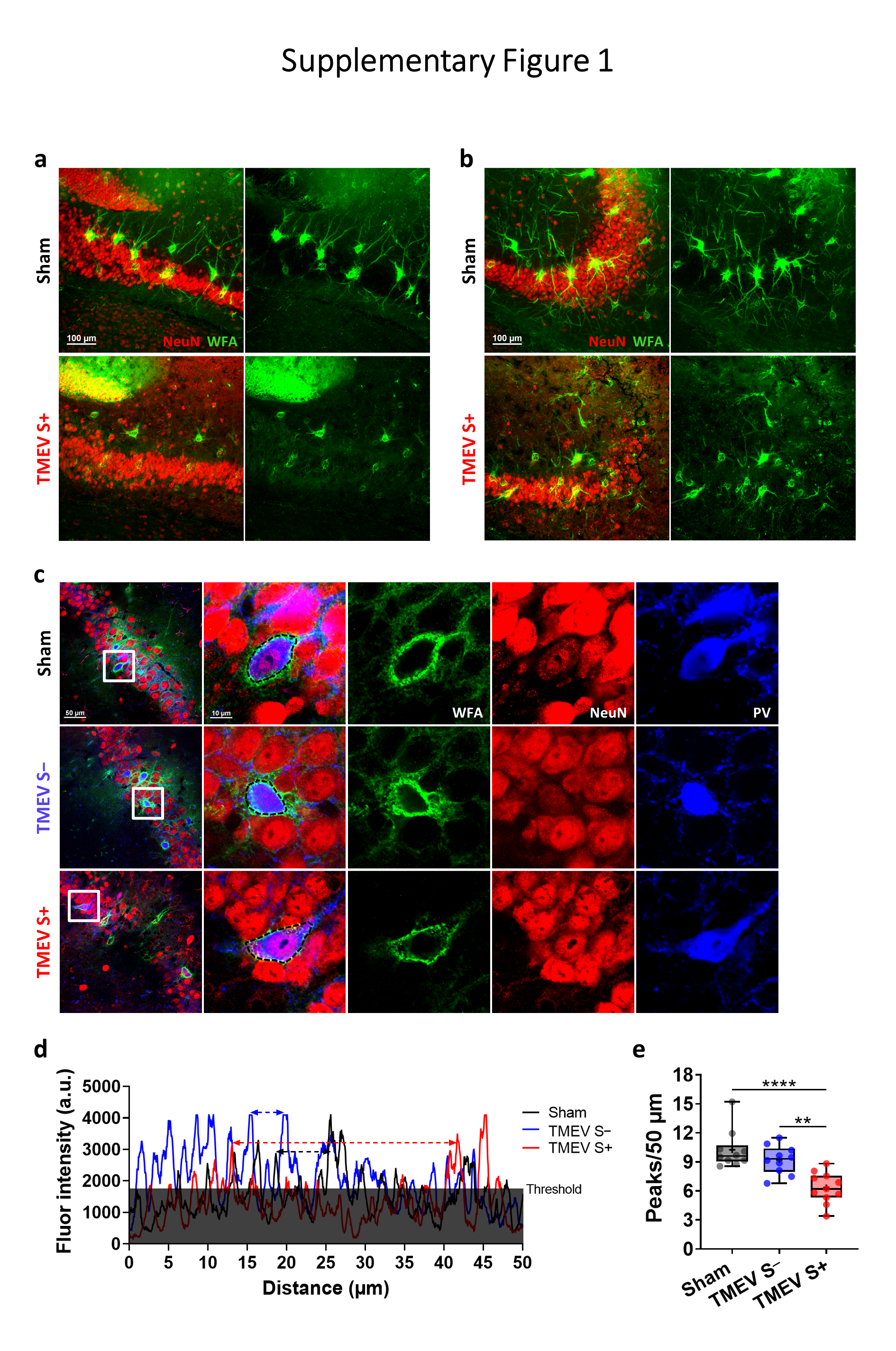

### Supplementary Fig 2

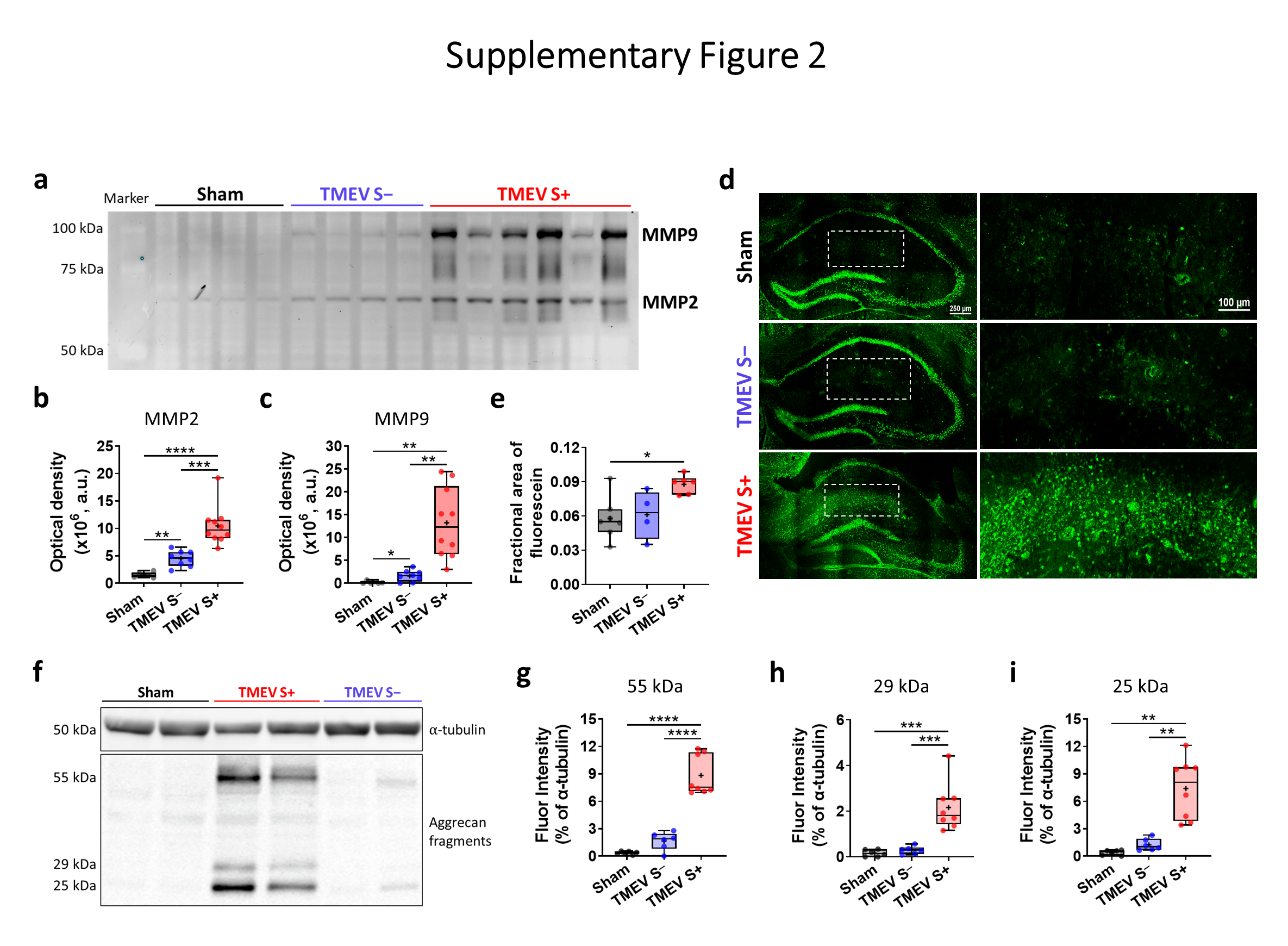

### Supplementary Fig 3

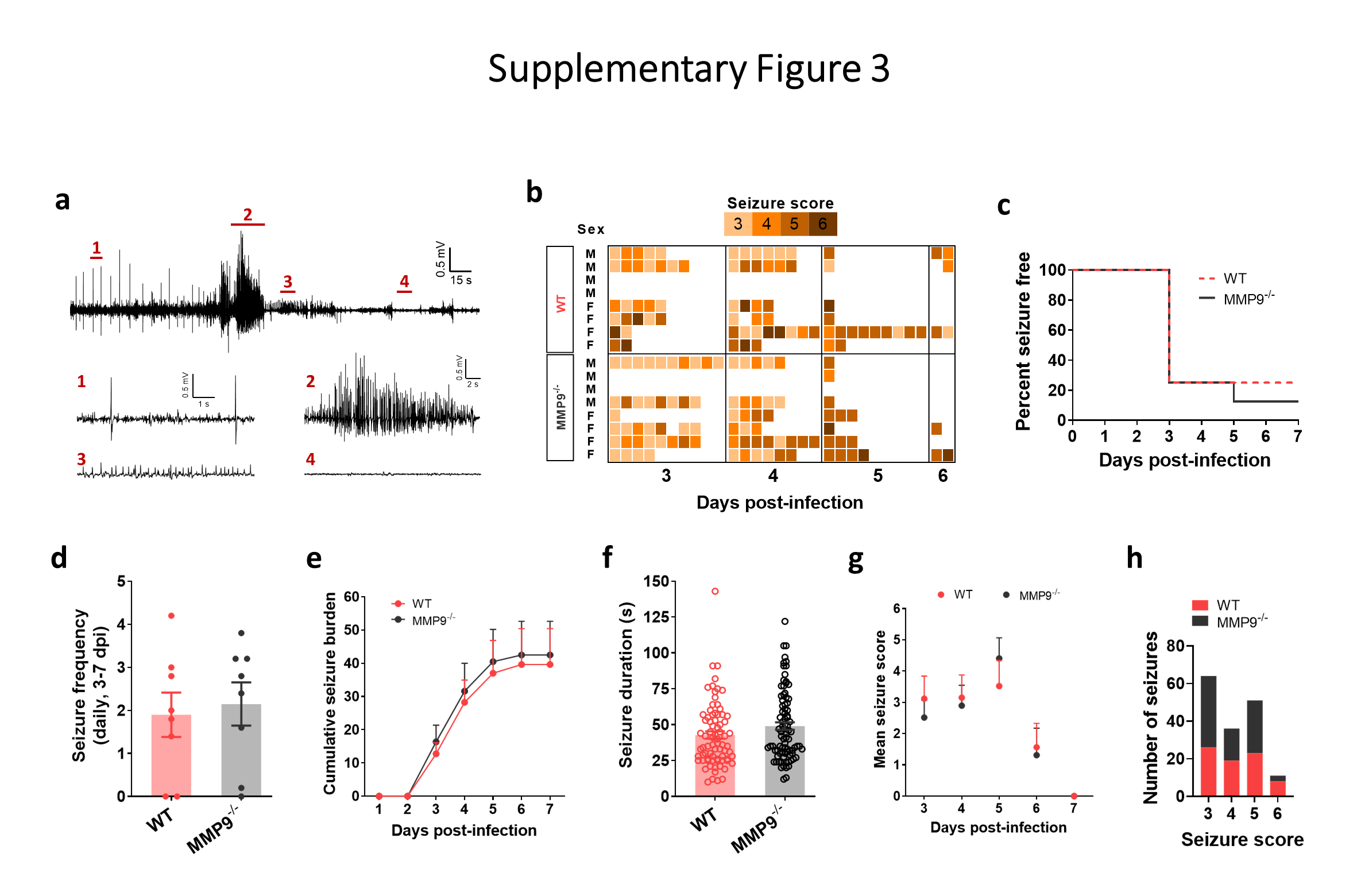

### Supplementary Fig 4

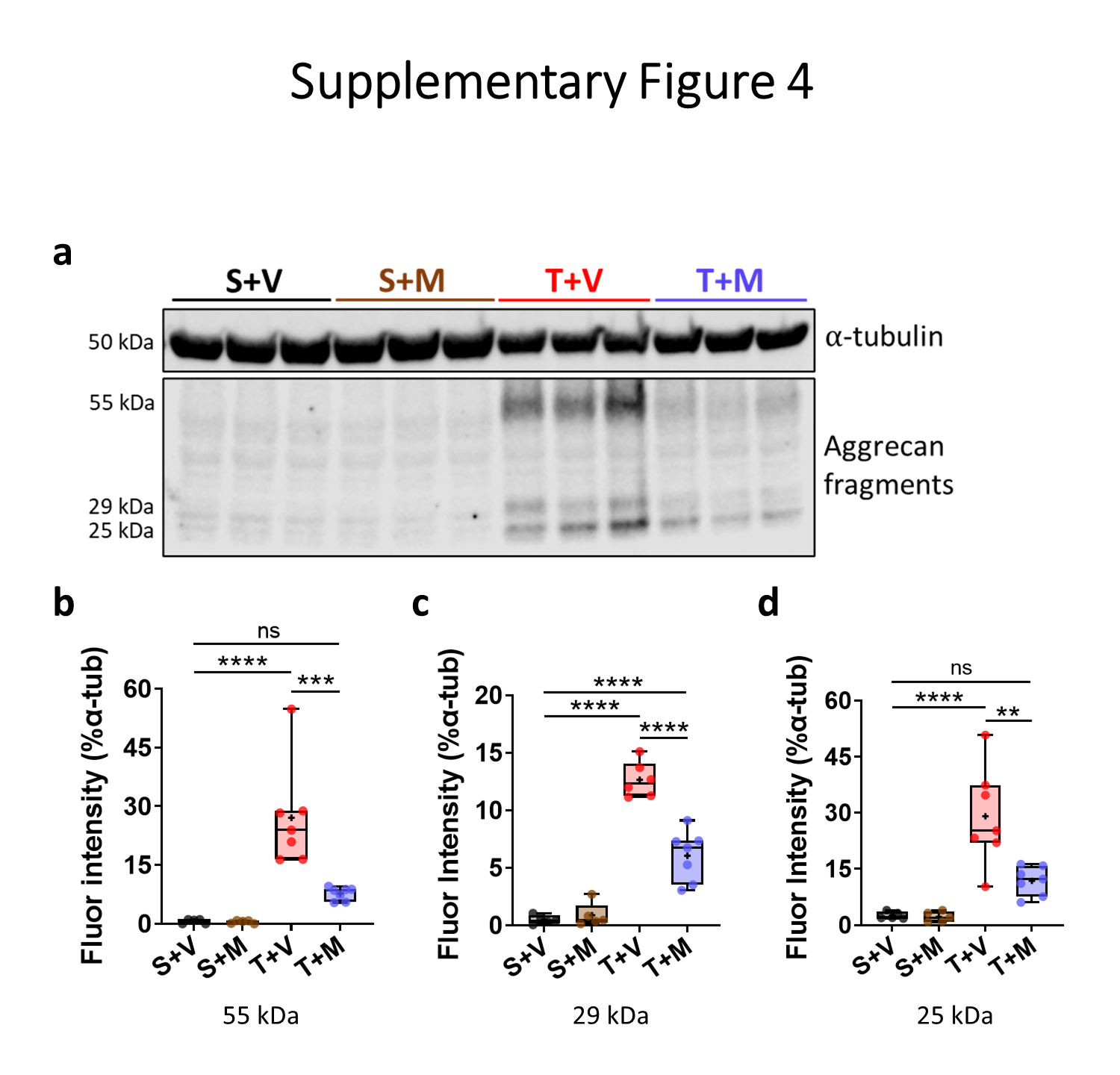

### Supplementary Fig 5

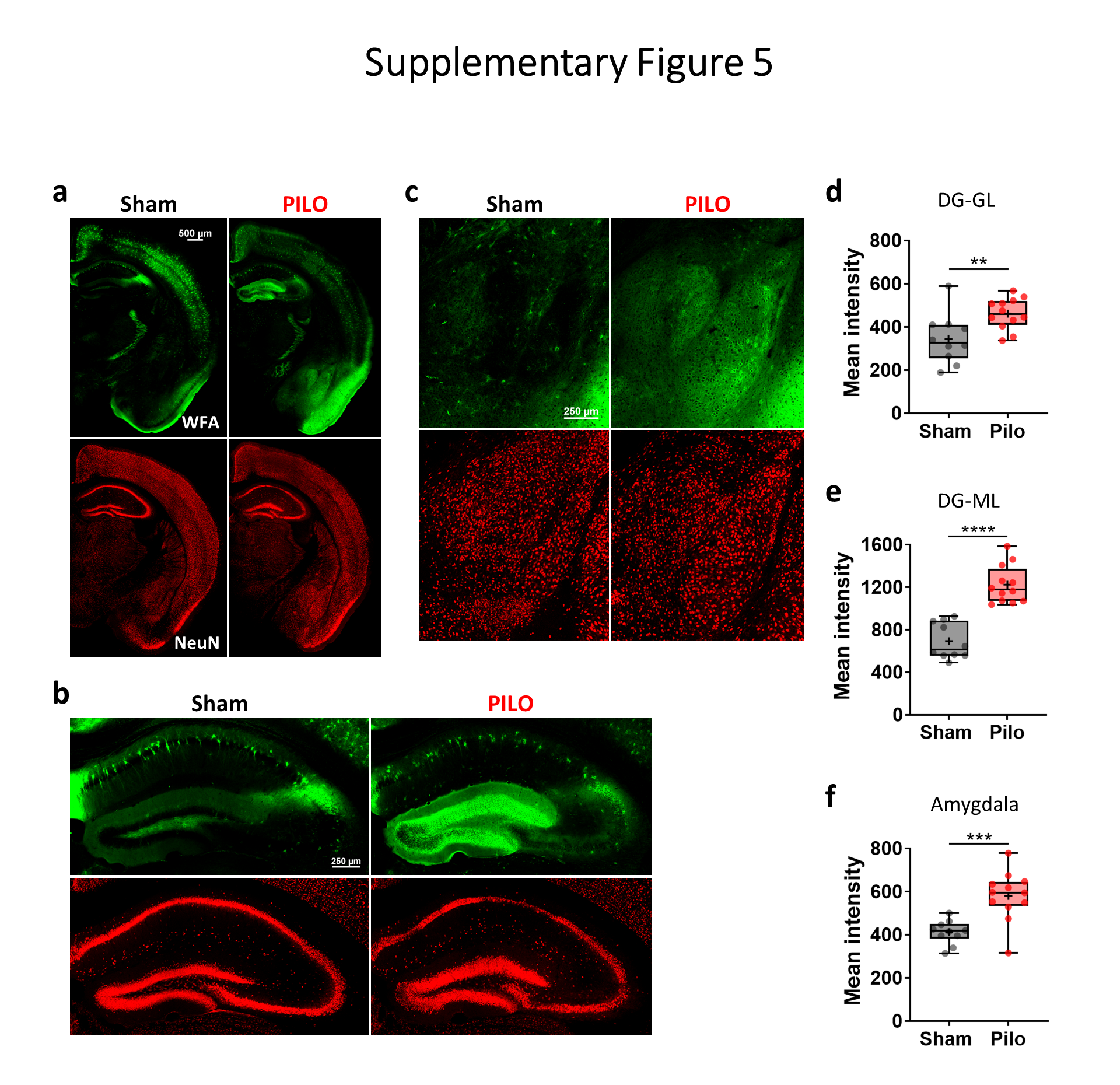

### Supplementary Fig 6

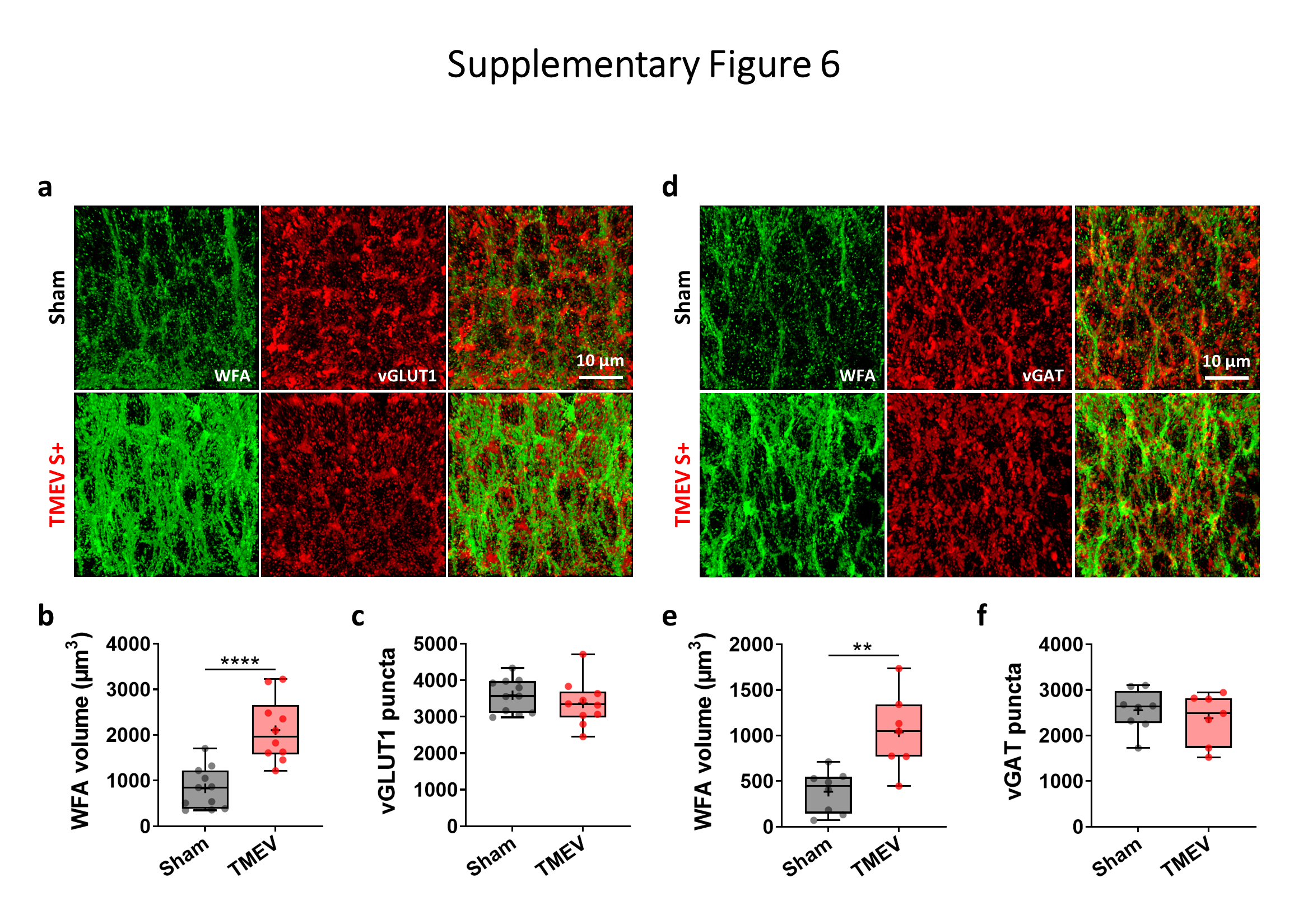

### Supplementary Fig 7

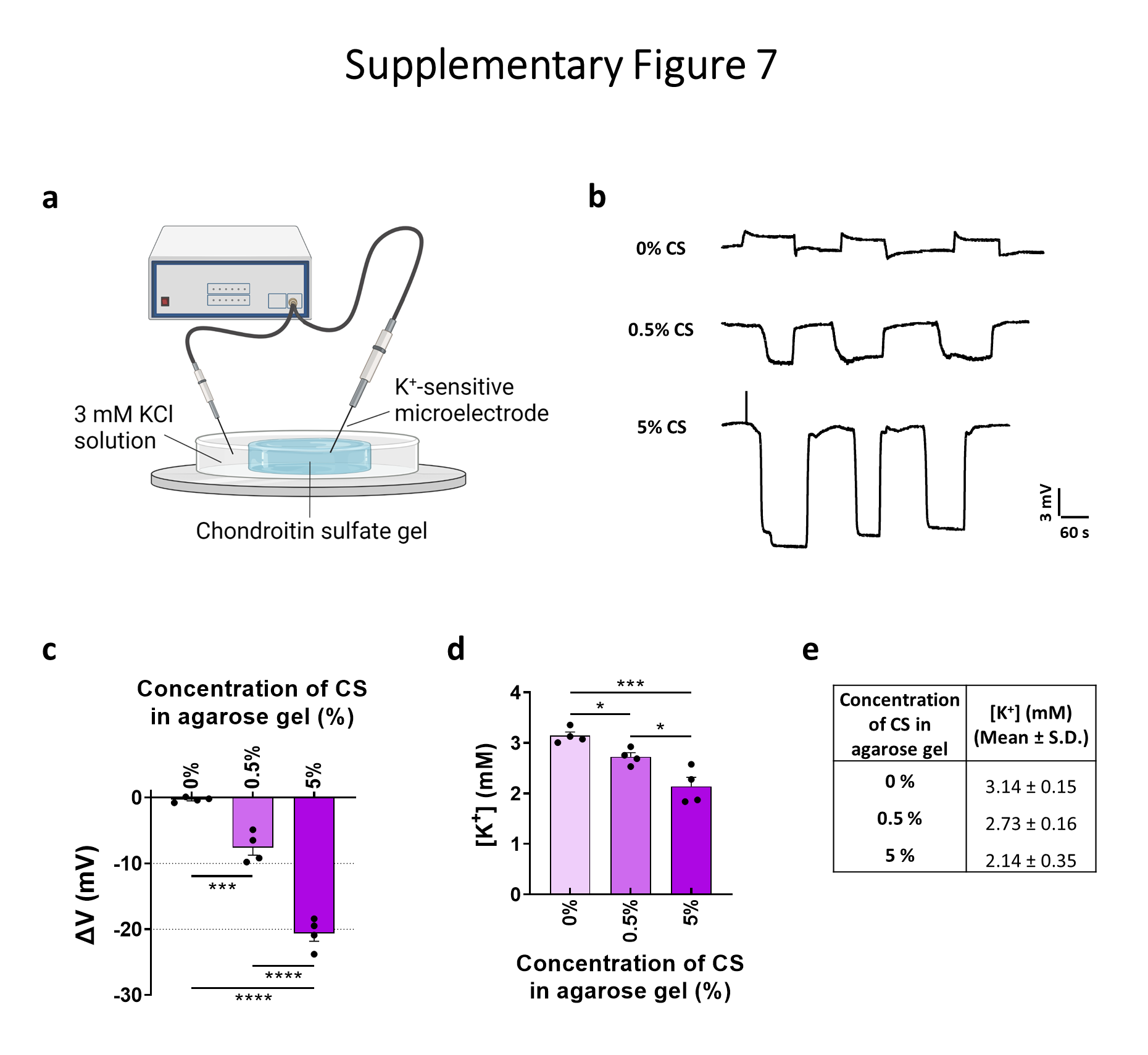
